# Integrated analysis of protein sequence and structure redefines viral diversity and the taxonomy of the *Flaviviridae*

**DOI:** 10.1101/2025.01.17.632993

**Authors:** Peter Simmonds, Anamarija Butković, Joe Grove, Richard Mayne, Jonathon C. O. Mifsud, Martin Beer, Jens Bukh, J. Felix Drexler, Amit Kapoor, Volker Lohmann, Donald B. Smith, Jack T. Stapleton, Nikos Vasilakis, Jens H. Kuhn

## Abstract

The *Flaviviridae* are a family of non-segmented positive-sense enveloped RNA viruses containing significant pathogens including hepatitis C virus and yellow fever virus. Recent large-scale metagenomic surveys have identified many diverse RNA viruses related to classical orthoflaviviruses and pestiviruses but quite different genome lengths and configurations, and with a hugely expanded host range that spans multiple animal phyla, including molluscs, cnidarians and stramenopiles,, and plants. Grouping of RNA-directed RNA polymerase (RdRP) hallmark gene sequences of flavivirus and ‘flavi-like’ viruses into four divergent clades and multiple lineages within them was congruent with helicase gene phylogeny, PPHMM profile comparisons, and comparison of RdRP protein structure predicted by AlphFold2. These results support their classification into the established order, *Amarillovirales*, in three families (*Flaviviridae, Pestiviridae*, and *Hepaciviridae*), and 14 genera. This taxonomic framework informed by RdRP hallmark gene evolutionary relationships provides a stable reference from which major genome re-organisational events can be understood.

## Introduction

The *Flaviviridae* are a family of positive-sense RNA viruses that incorporates several major human and veterinary pathogens, including hepatitis C virus (HCV) and a wide range of often highly virulent arthropod-borne viruses, including yellow fever virus (YFV) and dengue viruses^1^. Historically, the mosquito-borne YFV became the founding member of a growing group of ‘arthropod-borne viruses’, a term introduced in the 1940s^2^, although serologically split into two groups, A (Sindbis virus and relatives) and B (YFV and relatives)^3^. The taxonomic term ‘arbovirus’ was used in 1971 in the first report of the what became the International Committee on Taxonomy of Viruses (ICTV), with Arbovirus Group B renamed as the genus *Flavivirus* and group A as *Alphavirus* in the family *Togavivirdae.* This family also included the non-arbovirus transmitted genera *Rubivirus* (human rubella virus) and *Pestivirus* (bovine viral diarrhea virus (BVDV) and relatives infecting bovids and suids)^4^. Subsequent analyses of the growing amount of molecular, morphological, and serological data for these viruses indicated that the *Flavivirus* and *Pestivirus* genera should be removed from *Togaviridae* and re-classified into a new family, *Flaviviridae*^5,6^, a group that subsequently expanded to incorporate a third genus, *Hepacivirus*, for HCV^7^ and relatives in non-human primates, bats, and rodents (reviewed in^8^). Finally, a fourth genus, *Pegivirus*, was added in 2012 for a range of apparently non-pathogenic RNA viruses infecting humans (human pegivirus [HPgV]) and a broad range non-human primates, bats, and other mammas^9^ and one avian (goose) host^10^. Since then, there have been only minor changes to flavivirus classification, essentially limited to the expansion of the number of species assigned to the *Pestivirus, Pegivirus* and *Hepacivirus* genera^11^ and the renaming of the genus *Flavivirus* to *Orthoflavivirus*^11^.

Current members of the *Flaviviridae* have consistent genome organizations (positive-sense, strategies (synthesis of a single polyprotein with a conserved organization that is cleaved into structural proteins located at the N terminus and nonstructural proteins at the C terminus) (Fig. 1), and a primarily mammalian host range (in case of orthoflaviviruses, also arthropods)^1^. In addition to the RdRP gene, flaviviruses are homologous in their superfamily 2 helicase (NS3) and serine protease domain sequences^12^.

**Fig. 1.**
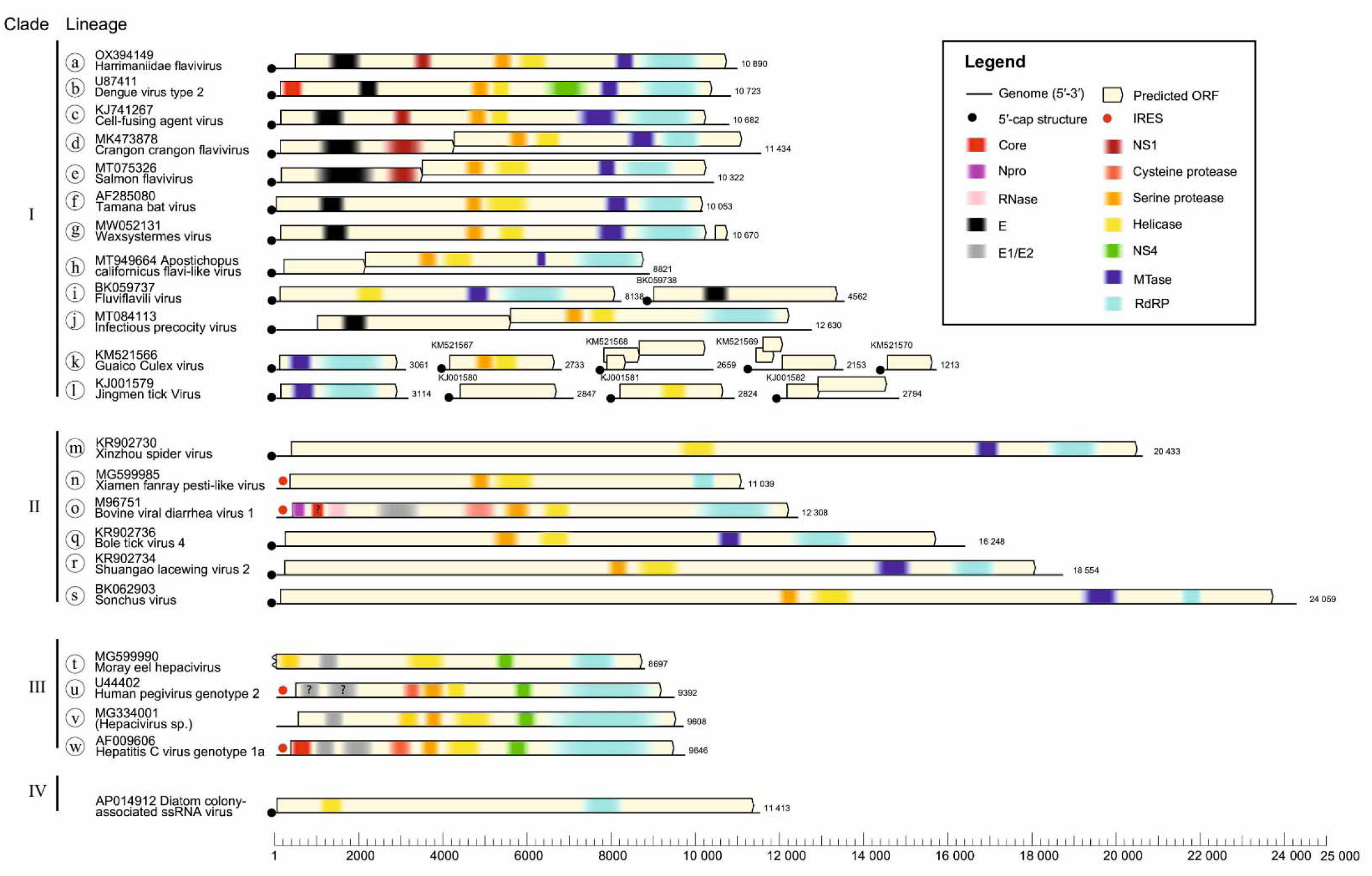
Organization of example genomes in each lineage of flaviviruses and ‘flavi-like’ viruses. Genome diagrams for the example viruses listed in Table 1 drawn to scale (lower scale bar) and main functional domains identified by InterProScan browser v. 103 (https://www.ebi.ac.uk/interpro/search/sequence/)^91^

**Table 1.**
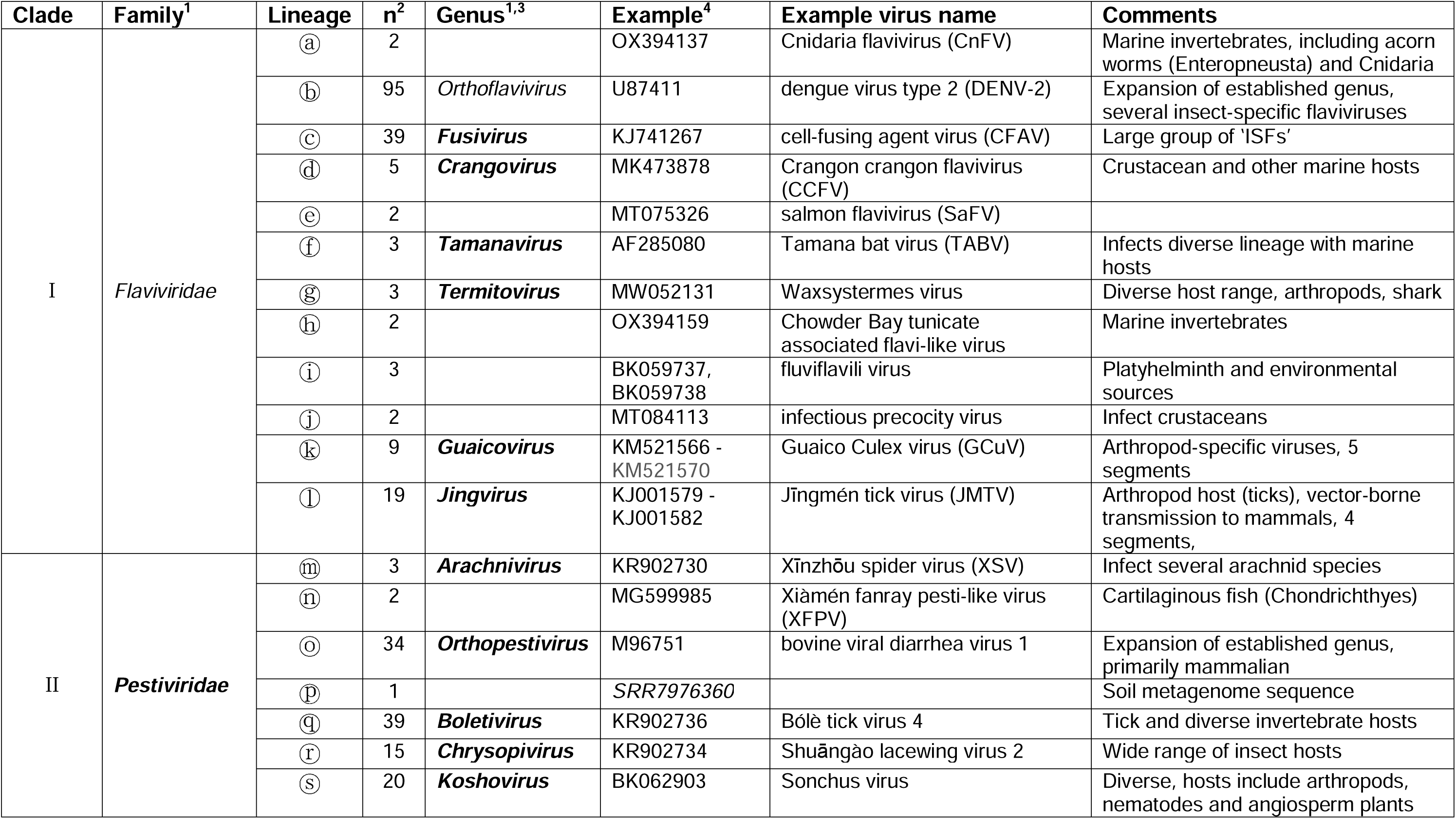
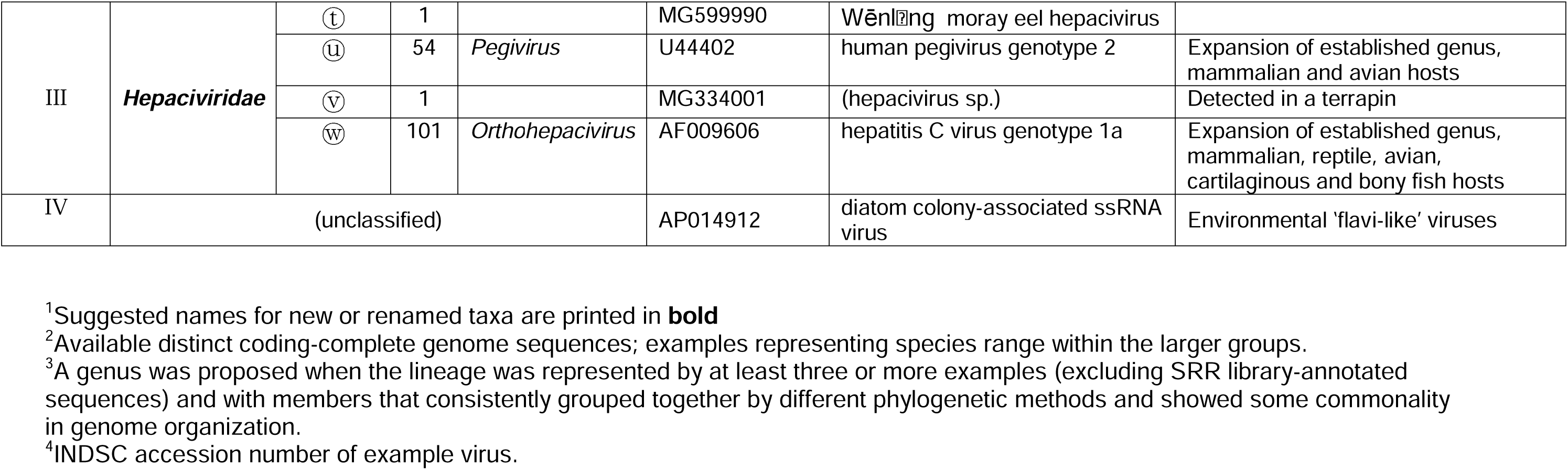
Listing of clades and lineages of flavivirus and ‘flavi-like’ viruses and suggested revised taxonomy.

However, within flaviviruses, there are also some major differences, including possession of structurally distinct capsid proteins (and the apparent lack of a capsid in pegiviruses), packaging mechanisms, polyprotein translation strategy that may be 5’-cap-dependent (orthoflaviviruses) or driven through an internal ribosomal entry site (IRES) in other flaviviruses (Fig. 1), and two disparate fusion glycoprotein systems. (Fig. 1). Indeed, other than the serine protease, helicase and RdRP, no nonstructural protein genes are homologous across viruses of all four genera^11^.

The more recent application of high throughput sequencing technologies to detect and genetically characterize viruses in a much wider range of potential hosts has transformed our understanding of flaviviral diversity and abundance^13^. Analyses particularly of arthropods and more recently fish and other aquatic life have created a case for a major expansion in viruses assigned to the existing genera, such as a range of novel insect-only (non-vectored) flaviviruses^14–19^ and the expansion of species assignments to the *Pestivirus*, *Pegivirus*, and *Hepacivirus* genera that infect primates, other mammals, and birds (reviewed in^8,20–23^).

More perplexing has been the discovery and characterization of a huge number of flavi-like viruses^24^, so-called because of their phylogenetic grouping of RdRP gene sequences with those of classified members of the *Flaviviridae*, but often possessing quite different genome organisations, genome lengths and host ranges from currently classified flaviviruses. For example, the so-called ‘large-genome flaviviruses (LGFs)’^13,25,26^ have genomes significantly longer than currently classified flaviviruses, the longest to date being maximus pesti-like virus’s genome of ≈40 kb^27^. Jīngmén tick virus (JMTV)^28^ and related jingmenviruses have multipartite genomes with four or more separate segments^29–32^. In addition, the majority of ‘flavi-like’ viruses have been discovered outside the primarily mammalian and vector host range of classified flaviviruses, being distributed across the animal kingdom, from poriferans (sponges)^27^, cnidarians (jellies)^33^, mollusks (squid)^34^, arthropods (insects^25,31,35^; diplurans^31^; scorpions^31^; crustaceans^33,34,36^), nematodes^37^, platyhelminths^38,39^) to echinoderms (sea cucumbers)^40^, hemichordates (acorn worms)^33^, cartilaginous and bony fish^13,33,41–44^, amphibians (frogs)^33,45^, and reptiles^434343^. Although only a few are known so far, ‘flavi-like’ viruses have also been discovered in stramenopiles (diatoms and oomycotes)^46,47^ and, remarkably, in angiosperm plants^30,48,49^.

These discoveries beg the question of how to classify these viruses and how to organize the current order *Amarillovirales* to best reflect their evolutionary relationships^13,50^. For example, by phylogenetic analysis of RdRP domain sequences, JMTV and other segmented ‘flavi-like’ viruses typically group closer to members of the *Orthoflavivirus* genus than to flaviviruses of other genera, but JMTV’s putative assignment to this genus conflicts with what had been previously considered family-defining characteristics across the *Flaviviridae* family, *i.e.*, a monopartite genome. The genomic diversity of ‘flavi-like’ viruses and expanded host ranges that now encompasses multiple kingdoms of eukaryotes profoundly questions how a virus family such as the *Flaviviridae* can be best defined^24^.

The future development of a robust taxonomy framework for flaviviruses, and indeed other virus families with similarly expanded genetic and genome organisational complexities, clearly requires a radically different set of criteria than those that have guided their current classification.

In this proposed update to the classification of flaviviruses, we have assigned primacy to the RdRP (hallmark) gene phylogeny using previously established principles for a genomics-based taxonomy of viruses^51^, and that assignments should be based on the most evolutionarily conserved gene within virus groups^52–54^. At present, most viruses are assigned into six realms that are divided based on their separate origins and possession of shared orthologous gene(s)^55–57^. Almost all RNA viruses, including flaviviruses, have been assigned to the realm *Riboviria* based on possession of evolutionarily related RdRPs that are distinct from those of cellular polymerases, substantiating their likely single origin. As the hallmark gene for ribovirians, its phylogenetic relationships therefore serve to best delineate the evolutionary history of RNA viruses. Re-casting virus taxonomy through evolutionary relationships of hallmark genes^51^ has furthermore enabled the development of a hierarchical higher-rank classification of RNA viruses, in which the family *Flaviviridae* has been assigned as a (single) member of the order, *Amarillovirales*, in the class *Flasuviricetes*, phylum *Kitrinoviricota*, and kingdom *Orthornavirae*.

This approach also acknowledges that the deeper evolution of all viruses may be punctuated by major genome reorganisations associated with changes in host range or ecologies. Examples of modular evolution in flaviviruses include multiple IRES exchanges within and between flaviviruses and viruses of the family, *Picornavirusae*^58,59^, and of acquisition of glycoproteins associated with commitment to vertebrate hosts^60^. Collectively, these create different evolutionary histories and phylogenetic relationships between different genome regions. While these evolutionary events can be represented in the form of reticulated trees or networks, the strictly hierarchical classification of viruses required by the ICTV (and indeed of other biological taxonomies) requires primacy to be assigned to hallmark genes in the creation of coherent taxonomy frameworks.

Accordingly, we present re-classification of flaviviruses and ‘flavi-like’ viruses grounded on varied analyses of RdRP hallmark gene relationships. This provides a robust, well-supported and coherent taxonomic framework that can serve as a scaffold for future expansion of this group of viruses. Moreover, our multi-modal approach may provide a template for re-examining other viral taxonomies, as we seek to organize the huge viral diversity revealed by meta-genomics.

## Results

To establish a new taxonomic framework for classified flaviviruses and unclassified ‘flavi-like’ viruses, we generated a dataset of representative sequences from currently classified members of the family *Flaviviridae* (viruses of 52, 19, 14, and 11 species in four genera, *Orthoflavivirus*, *Pestivirus*, *Hepacivirus* and *Pegivirus*, respectively;^1^) and a comprehensive sample of coding-complete sequences of currently unclassified ‘flavi-like’ viruses^60^ (listed in Table S1; Suppl. Data). This dataset included representatives from groups of segmented ‘flavi-like’ viruses, divergent ‘pesti-like’ viruses, many with extended genomes, many additional ‘hepaci-’ and ‘pegi-like’ viruses primarily found in marine vertebrates, plant-infecting ‘koshoviruses’ and a selection of ‘flavi-like’ viruses recovered from environmental samples such as diatom colony-associated virus (DCAV). Genome regions encoding the enzymatic domains of the RdRP and helicase genes were extracted for alignment and analysis.

### RNA-directed RNA polymerase domain (NS5/NS5B) amino-acid sequence phylogenies differentiate flaviviruses and ‘flavi-like’ viruses into four highly supported main clades

Our initial analysis focused on translated flavivirus and ‘flavi-like’ RdRP domain sequences. Using the distantly related tombusviruses as an outgroup, the RdRP domain phylogeny of our sequence set, calculated with IQ-TREE, resulted in four main bootstrap-supported clades (I–IV), each containing a number of bootstrap-supported lineages (labelled as Ia-l, IIm-s, IIIt-w anticlockwise around the tree) (Fig. 2). Viruses of the four currently established flavivirus genera were distributed in three of the four clades, with hepaciviruses and pegiviruses clustering together in and largely defining clade III. In this phylogeny, ‘jingmenviruses’, ‘tamanaviruses’, and numerous ‘insect-specific flaviviruses (ISFs)’ cluster with orthoflaviviruses in clade I. The viruses that were previously loosely defined as ‘LGFs’ join pestiviruses in clade II, and include diverse viruses distributed across multiple lineages, *eg*. Bólè tick virus (a potentially tick-vectored pathogen of mammals)^26^ in clade IIq and plant-infecting ‘koshoviruses’^30,48,49^, in clade IIs (Fig. 2; Table S1).

**Fig. 2.**
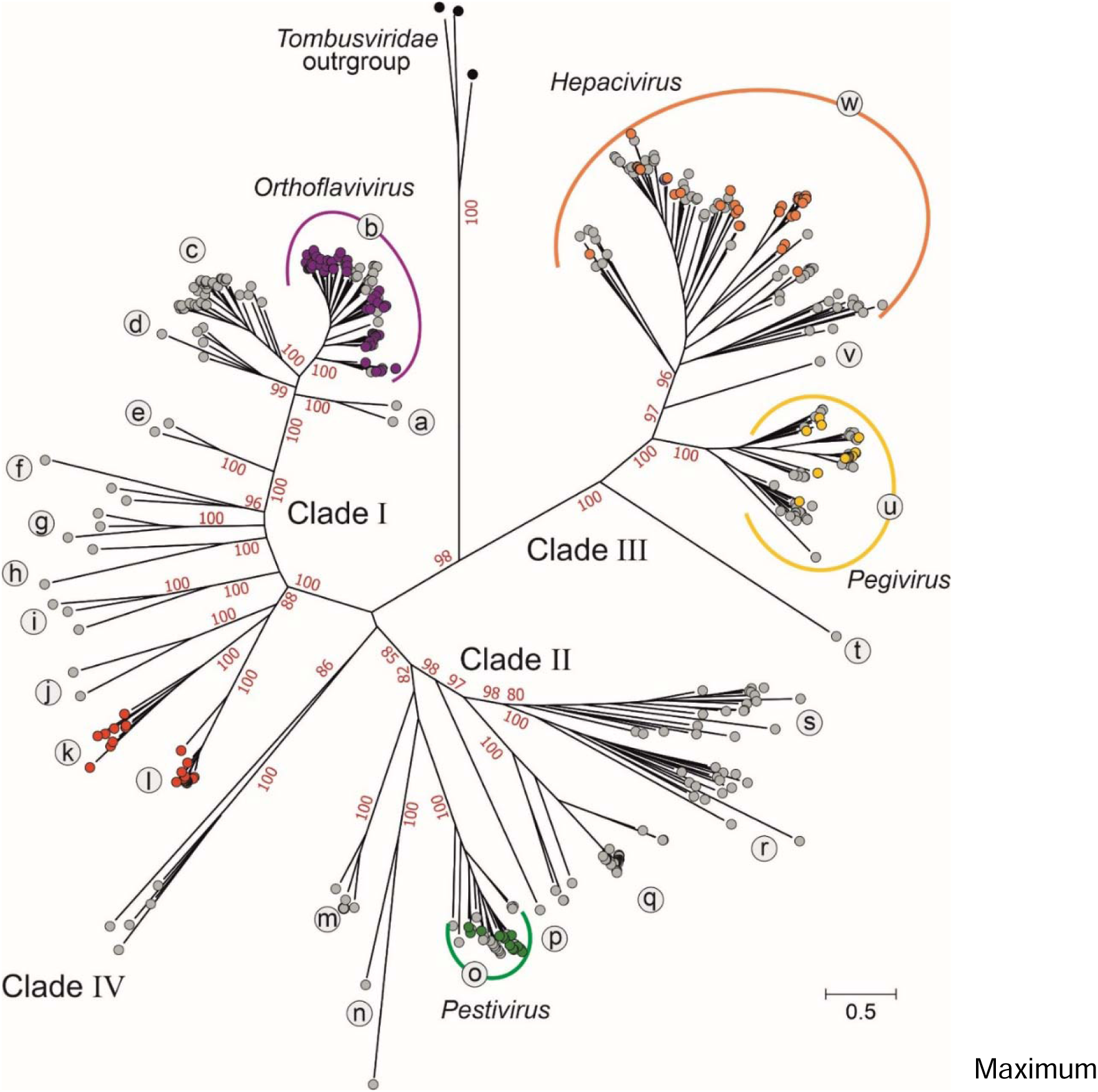
RNA-directed RNA polymerase domain (RdRP) amino-acid sequence phylogenies differentiate flaviviruses and ‘flavi-like’ viruses into four highly supported main clades. Maximum likelihood tree of aligned flavivirus and ‘flavi-like’ RNA-directed RNA polymerase (RdRP) domain amino-acid sequences, estimated using the LG+F+R10 model using IQ-TREE with tombusvirus sequences as an outgroup. Bootstrap support values for the main branches of the tree are shown in red if ≥70%. Already classified flaviviruses are shown in color-filled circles. The main clades are numbered I–IV and lineages are labeled with lower-case letters. The component sequences within each clade are provided in a fully annotated tree (RdRP_tree.NWK (File Sq; Suppl. Data) and listed in Table S1 (Suppl. data).

To investigate the robustness of the clade and lineage groupings, the IQ-TREE-based maximum-likelihood phylogeny generated for Fig. 2 was compared with trees generated through a temporal reconstruction using the Bayesian evolutionary analysis by sampling trees cross-platform program (BEAST) and using the protein distance-based unweighted pair group method with arithmetic mean (UPGMA) phylogeny method (Fig. 3). BEAST and UPGMA results reproduced clades I–IV with bootstrap support comparable to the original maximum likelihood analysis (Figs. 2, 3A). However, in the time-rooted BEAST tree, *Tombusviridae* became an inlier (Fig. 3C), in marked contrast to its clearly defined outgroup positions in the IQ-TREE and UPGMA trees. The grouping of individual sequences within lineages a–w defined by IQ-TREE analysis were almost entirely reproduced in the BEAST and UPGMA trees, with the exception of eight sequences that did not group with their lineages in the UPGMA tree (Fig. 3B). Despite these, the clade and lineage assignments were relatively robust to different evolutionary reconstruction methods.

**Fig. 3.**
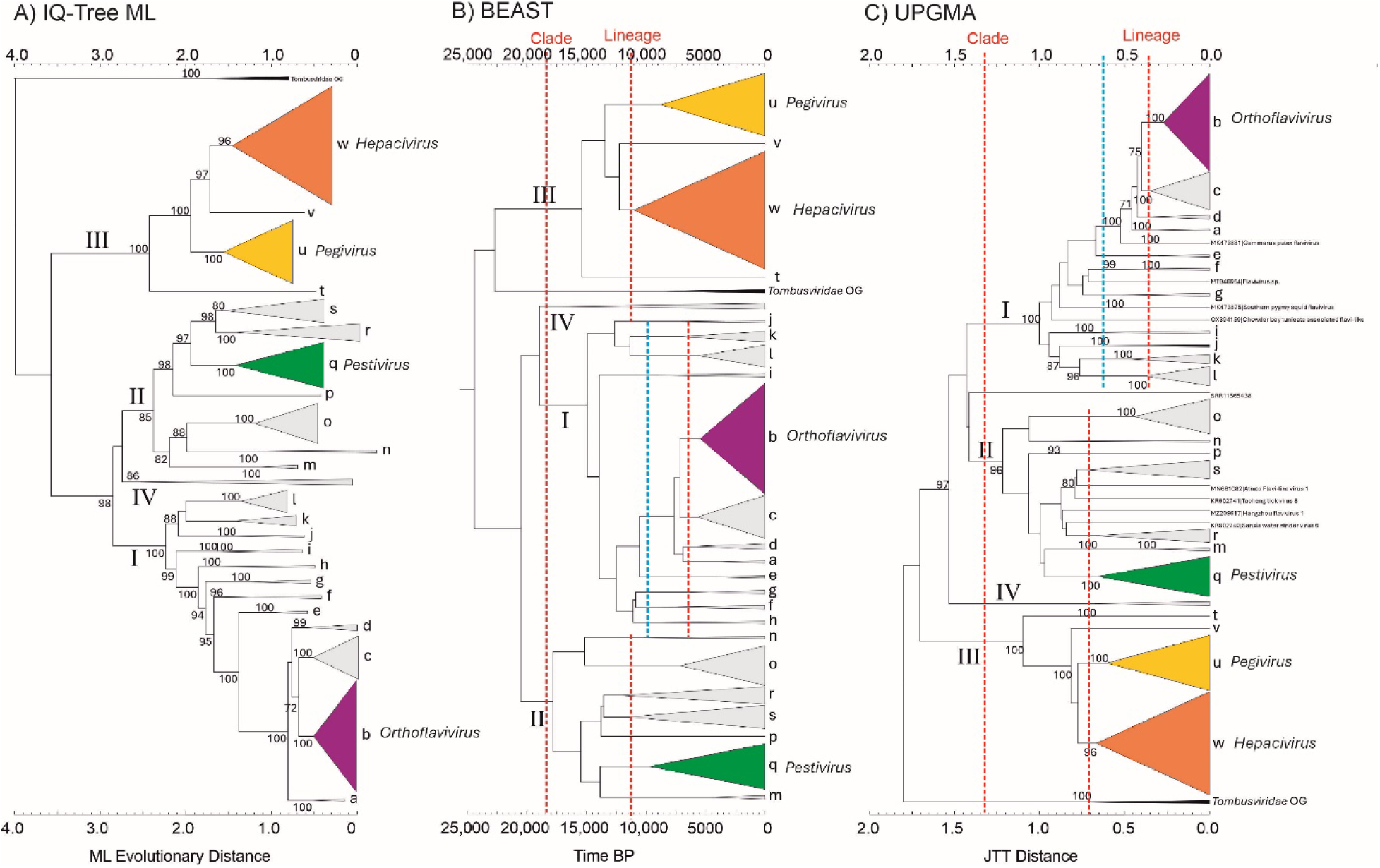
RNA-directed RNA polymerase domain (NS5/NS5B) amino-acid sequence phylogenies differentiate flaviviruses and ‘flavi-like’ viruses into four highly supported main clades. Phylogenetic trees constructed by likelihood (A, B) and distance-based (C) methods using flavivirus and ‘flavi-like’ RNA-directed RNA polymerase (RdRP) domain amino-acid sequences. Clades (I–IV) and lineages (a–w) labelled in each tree are based on those in Fig. 2. Tentative threshold levels of divergence separating clades and lineages are shown as red dotted lines in BEAST and UPGMA trees. An alternative threshold corresponding to the assignment of lineages Ia-Id to a common lineage is shown in a blue dotted line. Abbreviations: BP, before present; BEAST, Bayesian evolutionary analysis by sampling trees cross-platform program; JTT, Jones-Taylor-Thornton matrix; ML, maximum likelihood; UPGMA, unweighted pair group method with arithmetic mean.

These RdRP phylogenies provide an excellent framework for the taxonomic reorganization of the *Flaviviridae*. Nonetheless, we recognized that phylogenetic inference over these very large genetic distances is challenging and therefore sought to corroborate (or, indeed, refute) these apparent taxonomic groupings with alternative and complementary approaches.

### Helicase domain (NS3) amino-acid sequence phylogeny supports partition of flaviviruses and ‘flavi-like’ viruses into four main clades

While there is evidence for modular exchange of structural or assessor proteins among flaviviruses of the four established current genera, for instance acquisition of glycoproteins ^60,61^, it is less clear whether sequences encoding the core replication module of these viruses (including serine protease, helicase and RdRP) evolve as a unit or are similarly subject to genome exchange and rearrangements. To investigate this, and to complement our RdRP analyses, we deduced flavivirus and ‘flavi-like’ helicase domain amino acid sequences, aligned them, performed IQ-TREE analysis, and compared the result to the RdRP IQ-TREE tree using a tanglegram (Fig. 4). Phylogenetic groupings of viruses of the four current genera were highly concordant, with only minor differences in branching order within genera. The positions of ‘flavi-like’ viruses grouping with orthoflaviviruses were generally concordant between regions, but with some exceptions. For instance, there was a change in the topology of the deeper branches underlying the ‘LGF’, pestivirus and hepaci/pegivirus groupings, creating paraphyletic groups not observed in the RdRP tree. This result is not necessarily surprising because the helicase domain is shorter and more divergent than the RdRP domain and hence likely reflects lower resolution of relationships rather than indicating genome reorganization between the two domains. Other minor exceptions included Wēnl⍰ng moray eel hepacivirus (WMEHV) moving from an outlier position in clade III (lineage IIIt) into *Hepacivirus* in the helicase tree, a finding not incompatible with potential sequence errors in the deposited RdRP region sequence. Bólè tick virus 4 (BoTV4) fell within the IIq lineage in the RdRP domain tree, but as an outlier to pestiviruses in the helicase domain tree. Finally, ‘flavi-like’ viruses from environmental samples (clade IV in the RdRP tree) were located within the pestivirus-‘LGF’ branches in the helicase domain tree.

**Fig. 4.**
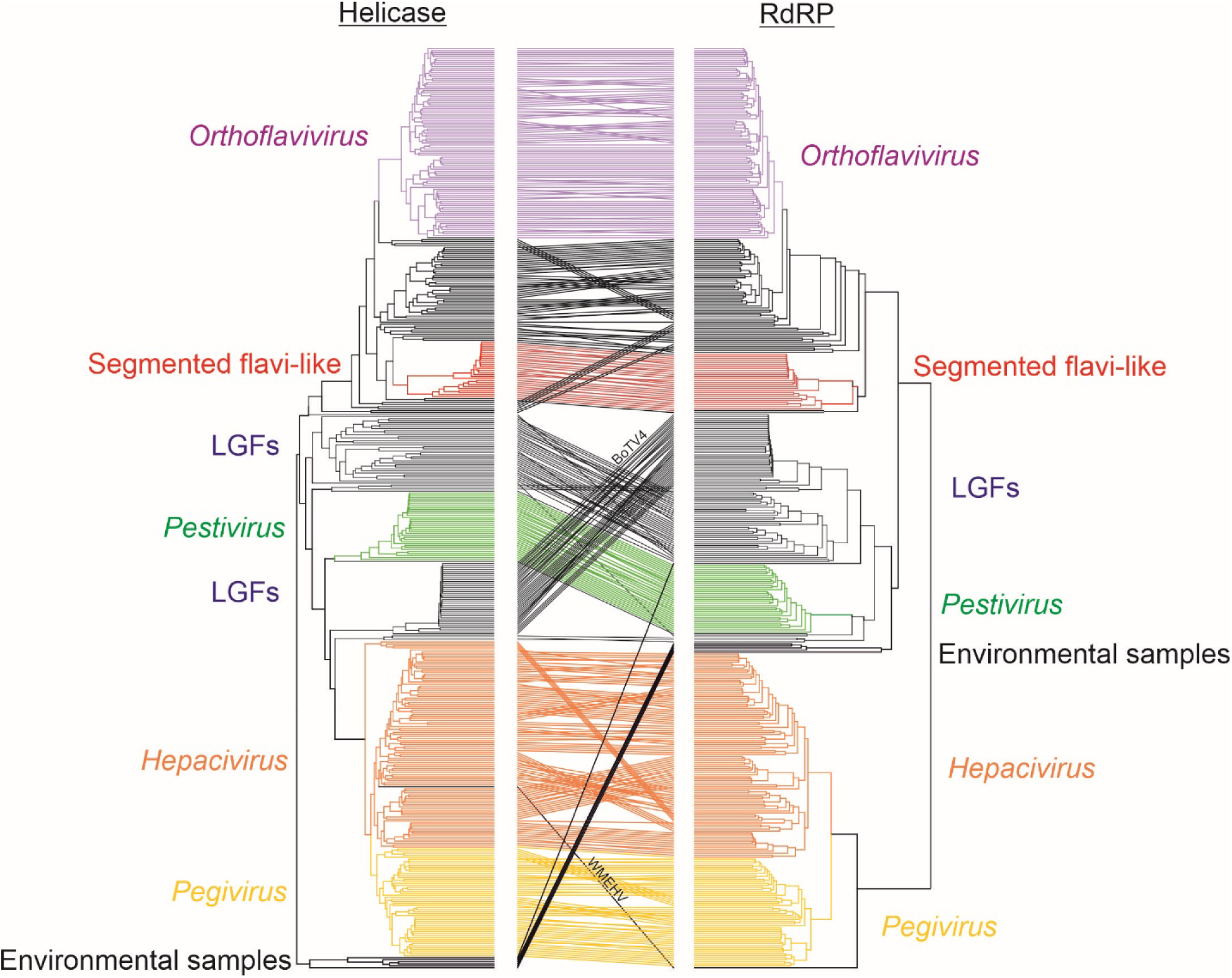
Helicase domain (NS3) amino-acid sequence phylogeny supports partition of flaviviruses and ‘flavi-like’ viruses into four main clades. Tanglegram of flavivirus and ‘flavi-like’ virus helicase and RNA-directed RNA polymerase (RdRP) domain sequences constructed from ML phylogenetic trees generated by IQ-TREE trees, with established flavivirus genera colored as in Figs. 2 and 3, and segmented ‘flavi-like’ viruses (lineages k and l in red). BoTV4, Bólè tick virus 4; LGFs, large genome flaviviruses, WMEHV, Wēnl⍰ng moray eel hepacivirus. A copy of the figure with the branches individually labelled is provided as Fig. S1.

### Structure-based comparisons of RNA-directed RNA polymerase (NS5/NS5B) recapitulate the sequence-based phylogeny

The function of any given protein is primarily a feature of its three-dimensional structure. Consequently, protein structure is fundamentally more conserved than the underlying protein sequence. The advent of accurate protein structure prediction through machine-learning (e.g., AlphaFold2) is enabling surveys of protein form and function at enormous scales^62,63^; and has driven the development of new high-throughput structure comparisons tools (e.g., Foldseek) that enable structure-guided inference of deep evolutionary relationships^64,65^.

In a recent investigation of glycoproteins, we systematically applied protein structure prediction to the *Flaviviridae*^60^, generating thousands of structures spanning the complete polyproteins of all viruses represented in the RdRP phylogeny (Fig. 2; Table S1, Suppl. Data). Drawing on this dataset, we generated complete NS5/NS5b RdRP domain structure predictions for each virus (see Methods). After filtering for prediction confidence and length, we analysed 400 RdRP structures using FoldTree^65^ to produce a structure-guided tree based on the local distance difference test (lddt) structural similarity metric. This structure-based approach does not explicitly consider protein sequence and, therefore, represents an independent recapitulation of the sequence-based phylogeny (Fig. 2).

The topology of the structure-based tree was remarkably similar to that of the sequence-based tree and supports the existence of the same four major clades (Fig. 5A). Moreover, most of the lineages represented in the sequence phylogeny were consistent in their position and composition. The only exceptions are lineages Ia, which in the structure-based tree formed a basal branch from lineage Ic, and lineages Ii and IIIv which moved subtly in relation to their neighboring clades. Clades Ij, IIn, and IIIt were lost from the structure-based tree due to filtering of lower confidence structural predictions. Major clade IV, containing ‘flavi-like’ viruses from environmental samples, also shifted position slightly, branching between clade II and the *Tombusviridae* outgroup in the structure-based tree. However, this divergent and basal taxon will likely remain difficult to place by any method without further discovery of similar viruses.

**Fig. 5.**
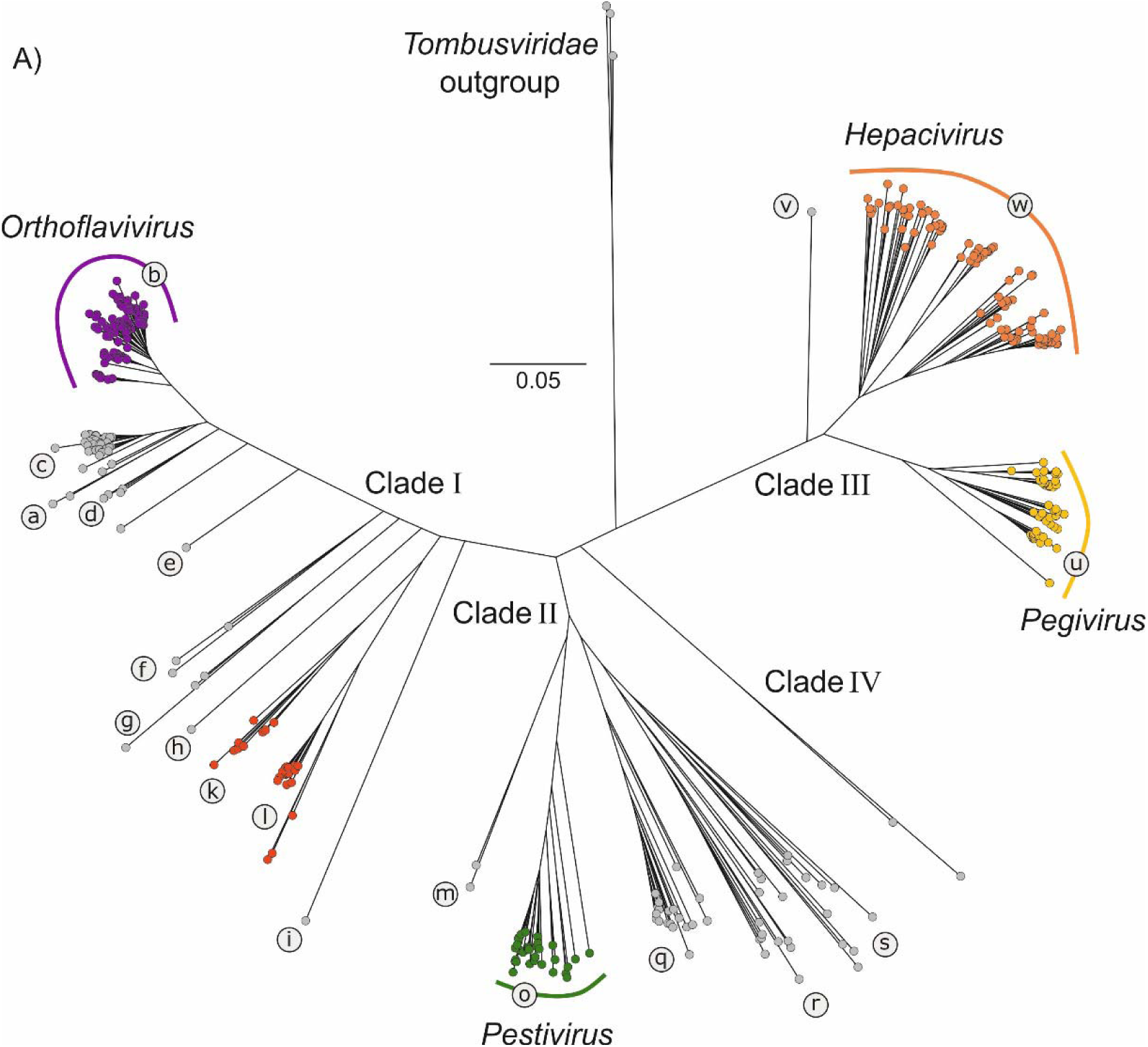
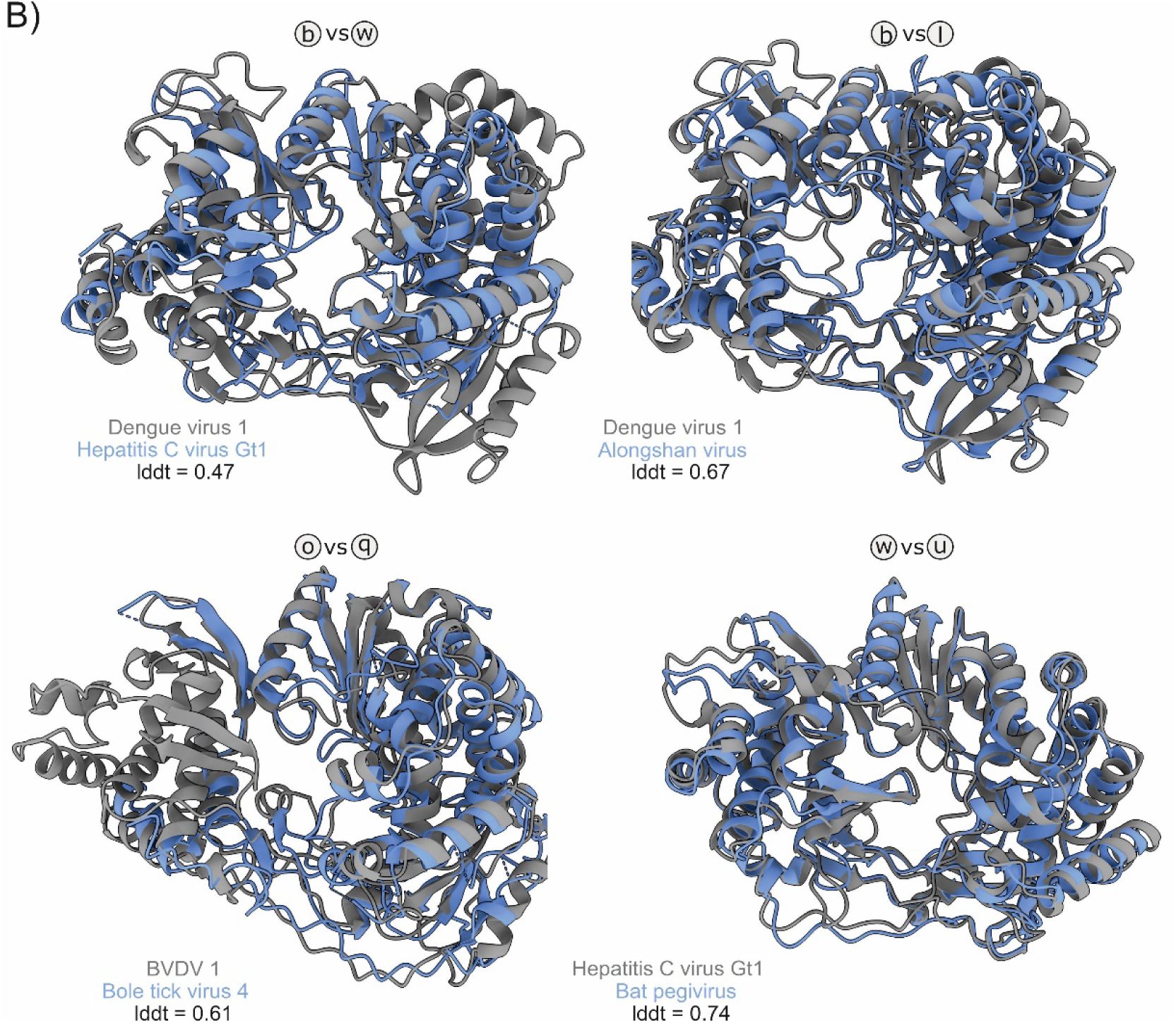
RNA-directed RNA polymerase domain (NS5/NS5B) structural comparison supports partition of flaviviruses and ‘flavi-like’ viruses into four main clades. A) Structure-based tree of 400 flaviviruses and ‘flavi-like’ viruses’ RdRP domains, derived from a lddt distance matrix (calculated by FoldTree^65^, powered by Foldseek^64^), scale bar indicates lddt distance (which is approximate to the inverse of the pairwise lddt score). Main clades and lineages are labelled as in Figure 2. B) Examples of aligned RdRP domain structures, color-coded as stated in the label, lineage identifiers (e.g. b vs w) indicate the position of the compared structures on the tree. lddt values represent structural similarity, with values of 1 being perfectly aligned identical structures. BVDV, bovine viral diarrhea virus.

Example structural alignments of clade representatives corroborate their distribution on the tree (Fig. 5B). Dengue virus 1 and HCV genotype 1 were at opposite ends of the tree (clade I and III) and their respective RdRPs align with a relatively low lddt score (0.47), whereas pairings within clade I, II and III give higher scores: 0.67, 0.61 and 0.74 respectively (note that an lddt score of 1.0 represents perfect alignment of identical structures). Thus, inferring evolutionary relatedness through structure-only analysis corroborates sequence-based approaches and is highly supportive of the organization of flaviviruses and ‘flavi-like’ viruses into four main clades.

### Alignment-free hidden Markov model homology analysis supports partition of flaviviruses and ‘flavi-like’ viruses into four main clades

Genome Relationships Applied to Virus Taxonomy (GRAViTy) is a non-supervised, alignment free method to assess the relatedness of virus genome sequences though calculation of protein profile hidden Markov model (PPHMM) homologies and through metrics of genome organization such as the order and orientation of genes^66^. We performed GRAViTy analysis using our flavivirus and ‘flavi-like’ virus dataset (Table S1) for phylogeny and RdRP structure comparisons; genomes of segmented ‘jingmenviruses’ were concatenated in order of segment length from short-to-long. Results were remarkably concordant with those determined by RdRP and helicase domain phylogenies (Figs. 2–4), with bootstrap-supported segregation of the same sequences into three main clades I–III (Fig. 6A). ‘Flavi-like’ viruses from environmental samples (formed an outlier position as clade IV.

**Fig. 6.**
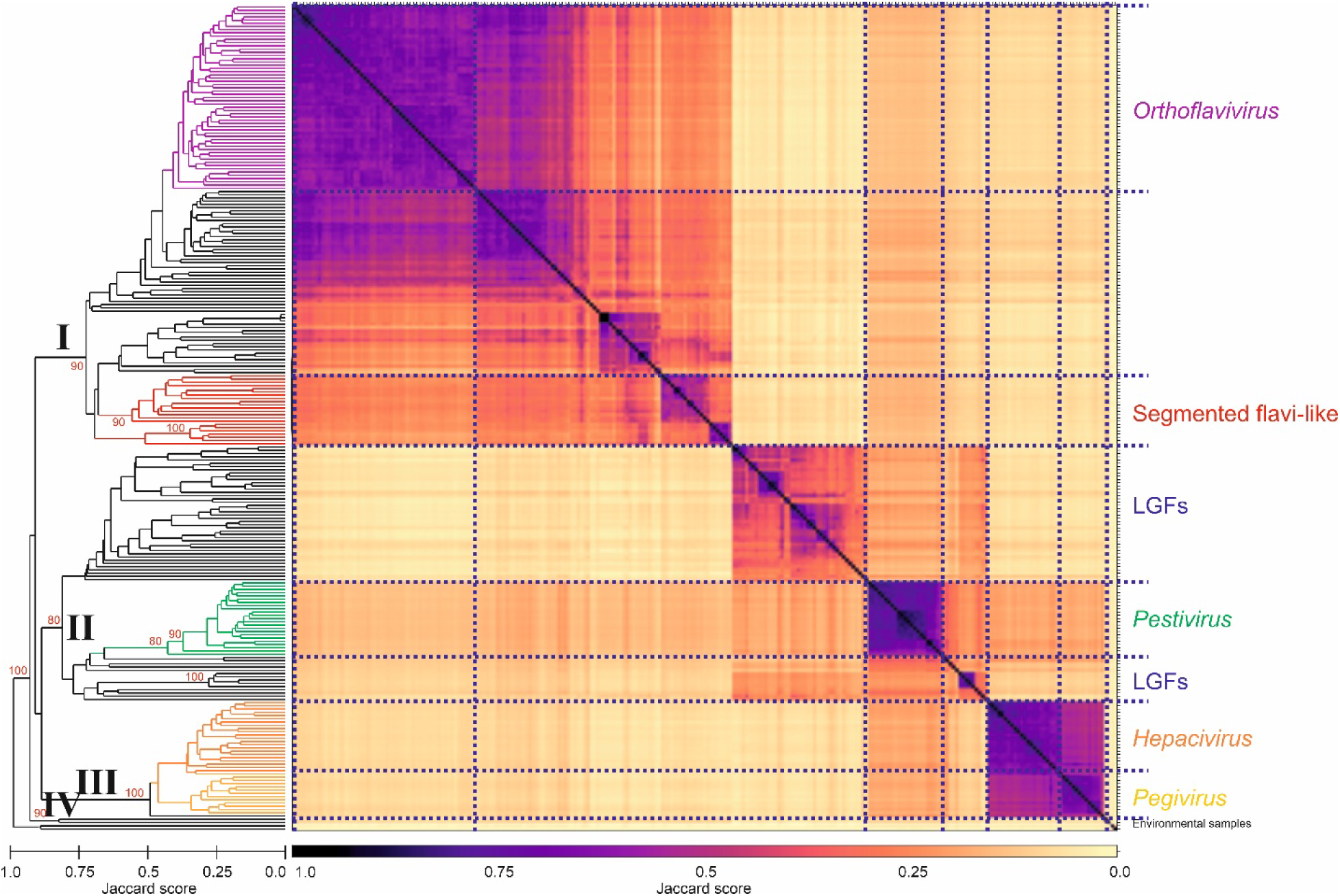
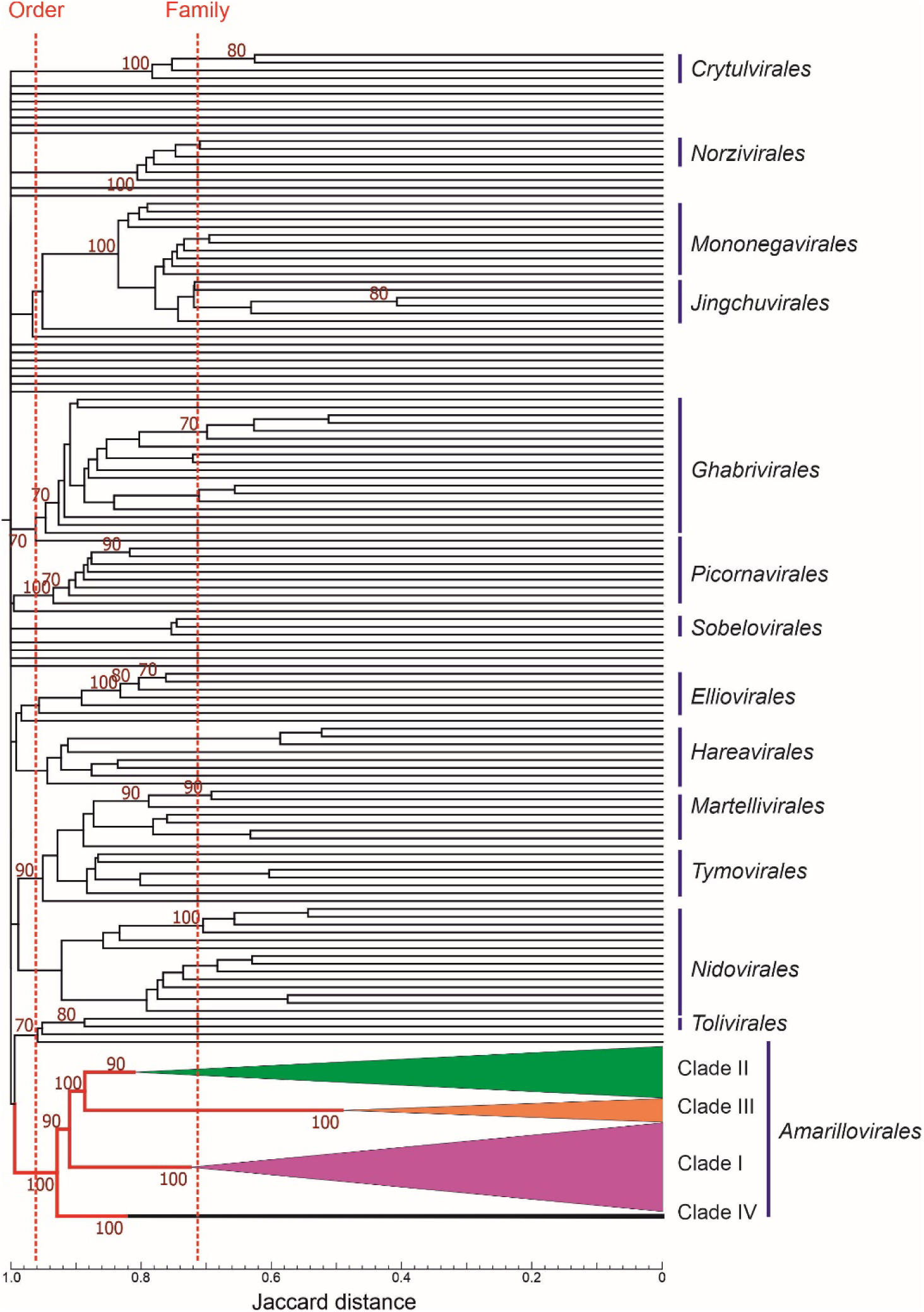
Alignment-free hidden Markov model homology analysis supports partition of flaviviruses and ‘flavi-like’ viruses into one order and four family rank clades. A)GRAViTy Jaccard distances calculated for classified flaviviruses and ‘flavi-like viruses and a representative member of each established RNA virus family in ribovirian kingdom *Orthornavirae* (n=135), showing approximate demarcation thresholds for orders and families (dashed vertical red lines). Bootstrap support values (10 iterations) are shown in red if ≥70%. A copy of the figure with the branches individually labelled is provided as Fig. S3. B) Heatmap and dendrogram depicting relationships among classified flaviviruses and ‘flavi-like’ viruses. Clades I–IV identified in the RdRP phylogeny (Fig. 2) were added to equivalent branches in dendrogram. Bootstrap support values (10 iterations) for deeper branches are shown in red if ≥70%. A copy of the figure with the branches individually labelled is provided as Fig. S2.s

To depict the wider inter-relationships of classified flaviviruses and ‘flavi-like’ viruses, we compared their sequences with sequences representing viruses of all other established ribovirian families. Flaviviruses and ‘flavi-like’ viruses were monophyletic in the GRAViTy dendrogram, supporting their assignment to a common higher taxonomic rank, the established order *Amarillovirales* (Fig. 6B). The Jaccard distances calculated by GRAViTy do not provide a precise quantitative estimate of evolutionary distance or thresholds for taxonomic assignments. However, ribovirian families differ from each other in the distance range of 0.7–0.85, whereas viruses within families typically are associated with Jaccard distances of approximately 0.9 (Fig. 6B). Although very general, the distances between clades I–IV were within the range of between-family distances elsewhere in the dendrogram, whereas their combined grouping occurs at the level of order assignments for other virus families assigned to orders (e.g., *Nidovirales* and *Picornavirales*).

### Taxonomic groupings correlate with genome properties and host range

Within clade I, lineage Ib contains the currently classified members of the *Orthoflavirus* genus, as well as a high number of currently unclassified primarily insect-specific flaviviruses (ISFs). Lineage Ic is similarly populated by ISFs, including the current unclassified cell fusing agent virus (CFAV), consistent with a previous proposed assignment to a new genus within the *Flaviviridae*^67,64,66^. The similarly divergent Tamana bat virus (TABV) falls in the very diverse lineage If, that also contains ‘flavi-like’ viruses infecting lumpfish (*Cyclopterus_lumpus*) and the pygmy squid (*Xipholeptos notoides*). The segmented JMTV and Guaico Culex virus (GCuV) and their relatives have been assigned to lineages Ik and Il, adjacent to lineage Ij containing the non-segmented infectious precocity virus. Extending the host range of this clade were Cnidaria flavivirus and Harrimaniidae flavivirus infecting basal metazoa such as jellies and acorn worms^33^ in lineage Ia. Despite the evident lineage (and host) diversity, clade I is clearly monophyletic with a bootstrap-supported long branch separating members from other clades.

Clade II similarly sub-divides into a number of bootstrap-supported lineages IIm – IIs, with currently classified pestiviruses clustering exclusively in lineage IIo along with currently unassigned more divergent viruses exclusively infecting vertebrates, as does lineage IIn (in fish). Viruses assigned to other lineages within clade II are arthropod-hosted with the striking exception of the plant-infecting Apis flavivirus, carrot flavi-like virus 1, Gentian Kobu-sho-associated virus, Coptis virus 1 and Sonchus virus 1 that form a sub-group, termed ‘koshoviruses’, within lineage IIs^48,49,68^. Arthropod and plant-infecting members of clade II frequently showed substantially longer genomes than the vertebrate-infecting members of lineages IIn and IIo (ranges 11038–11450 and 11555–15154 respectively) compared with lineage IIm (20432–22622), IIq (13599–18696), IIr (14731–26314) and IIs (18749–27708).

Members of clade III were all non-segmented, possessed similar genome lengths (8294–12290) and formed a well-defined separate grouping from other flaviviruses. In contrast to the wide host range of clade I and II, all clade III members had presumed or demonstrated vertebrate hosts, spanning a wide range of mammals, birds, reptiles, and bony and cartilaginous fish. Two lineages, IIIu and IIIw, contained currently classified members of the *Pegivirus* and *Hepacivirus* genera along with a range of more divergent viruses with a greater host range beyond mammals and birds.

### Genomic relationships between flaviviruses and ‘flavi-like’ viruses provides a basis for their classification

The reproducible phylogenetic relationships between flaviviruses and ‘flavi-like’ viruses using different tree construction methods in the RdRP domain sequences (Fig. 2 & 3) were recapitulated by protein structure-guided analysis (Fig. 5). RdRP is evidently co-evolving with the helicase gene (Fig. 4), suggesting that the essential replicase of flaviviruses and ‘flavi-like’ viruses traces a single coherent evolutionary history. This is also reflected in the relationships identified by an alignment-free method for analyzing whole genome sequences (Fig. 6). Therefore, we are confident that the foundational RdRP phylogeny in Figure 2 provides a robust framework for a genomics-based re-classification of flaviviruses.

These analyses based on clade and lineage relationships provide a framework that might map clades and lineages onto families and genera. However, there are quite variable branch lengths within lineages (Fig. 2) and differences in thresholds that split lineages in BEAST and UPGMA tree across clades I, II and III (Fig. 3). These observations indicate that a purely cladistic classification may not conform to specific sequence divergence thresholds that are often used elsewhere in virus taxonomy. Indeed, formal comparison of mean amino acid sequence identities between and within lineages in the three clades showed considerable variation (Fig. S4; Suppl. Data), particularly in clade I where sequence identities between lineages Ia, Ib, Ic, and Id were all much greater than within-group values of other lineages (notably If, Ih, Ii and Ij, and lineage n in clade II). For classification purposes, it might be considered that these four lineages were assigned as a group equivalent to those formed in other lineages; this would create a higher threshold in the UPGMA and BEAST trees (blue dotted line in Figs. 3B and 3C) that is more consistent with between lineage thresholds in clades II and III.

With this caveat, we can make some tentative proposals for the re-classification of flaviviruses that accommodates the large number of additional ‘flavi-like’ viruses described since the last classification of the family^1^ (Table 1). The following findings were considered in the design of a new flavivirus taxonomy:

A. Flavivirus and ‘flavi-like’ viruses form a monophyletic group in comparison with all other members of the *Riboviria*, enabling their assignment to a single taxonomic rank. Although not precise, divergence among flaviviruses at this rank was comparable to that of members of virus orders in GRAViTy analysis (Fig. 5B). We therefore propose that all flavivirus and ‘flavi-like’ viruses can be assigned to the established order *Amarillovirales*.
B. The level of sequence divergence among members of clades I–IV was comparable to inter-family distances of other RNA viruses on GRAViTy analysis, and we therefore suggest assignment of members of lineages I, II, and III to the new families. We provisionally name these *Flaviviridae*, ‘*Pestiviridae*’, and ‘*Hepaciviridae*’, reflecting the historical key virus members of these families.
C. While a separate, bootstrap-supported lineage IV was consistently observed, its members are highly divergent genetically and derive from environmental samples. Where hosts are suspected, these are extremely diverse and require further verification. We therefore do not suggesting creating a family for lineage IV at this time.
D. All three suggested families contain a number of bootstrap-supported clades of sequences, often with distinct genome organisations, lengths and host tropisms. We suggest that the clade assignments can be used for genus demarcation in certain circumstances:

a. Clades Ib, IIo, IIIu, and IIIw contain currently classified flaviviruses and should be assigned in the revised taxonomy. We suggest that the current genus names *Orthoflavivirus*, *Pegivirus*, and *Hepacivirus* are retained, with *Pestivirus* modified to ‘*Orthopestivirus*’ (because of the proposed *Pestiviridae* family name) and *Hepacivirus* modified to ‘*Orthohepacivirus*’ (because of the proposed *Hepaciviridae* family name) to maintain continuity with established nomenclature while acknowledging the greatly expanded diversity of viruses assigned to each.
b. Criteria for designating further genera include:

i. Bootstrap-supported groupings consistent with multiple methods
ii. A minimum of three members so that the extent of grouping can be assessed
iii. Priority should be given to clades containing previously described and well-characterised viruses, such as cell fusing agent virus (CFAV) and Tamana bat virus (TABV).
c. On this basis, we propose the creation and naming of a total of 10 additional genera in addition to the four currently assigned (Table 1).
d. An alternative classification within clade I would assign members of lineages Ia, Ib, Ic, and Id to the same genus to provide greater comparability with genetic divergence thresholds between genera elsewhere. The name *Orthoflavivirus* could then be applied to a greatly expanded range of viruses including CFAV, while its component lineages could be assigned as the subgenera ‘*Euflavivirus*’ (true flaviviruses), ‘*Crangovirus*’, and *iFusivirus*’.

## Discussion

The RdRP domain is considered the “hallmark gene” for the classification of RNA viruses, primarily because comparison of RdRP sequences provided evidence for their monophyletic origin separate from all known cellular polymerases^56,69^. Using the RdRP phylogeny as the reference point, the evolutionary history of flaviviruses and ‘flavi-like’ viruses has evidently been punctuated by multiple genome re-organizations, expansions, exchange of structural sequence modules, and genome segmentation^13,50,60^. The continuous discovery of ‘flavi-like’ viruses in highly divergent hosts means that it is no longer possible to use the criteria of genome length, coding strategy and host range that had been used for taxonomic placement of currently classified flaviviruses.

While the evolutionary history of these virus could be more completely and better represented as a multi-dimensional network or reticulate tree, the requirement of all biological taxonomies for a hierarchical classification necessitates selection of a common marker gene present in all clades. Thus, the confirmed grouping of already classified flaviviruses and ‘flavi-like’ viruses into established order *Amarillovirales* and their suggested assignment to three families and 14 genera (Table 1) is based on the phylogeny of RdRP. Other genome regions exhibit much more horizontal gene transfer, resulting in distinct evolutionary histories that may even originate from cellular life (e.g., E^rns^, found in the pestivirus and ‘LGF’, is of bacterial origin^60^); clearly, organization based on these genomic regions would confound taxonomic classification.

We observed a primary division into four clades, all of which are bootstrap-supported by each of the methods we used. The established genus assignments were replicated in these analyses, but there is a much closer relationship of hepaciviruses and pegiviruses than between or to the other genera. Indeed, hepaciviruses and pegiviruses cluster in one main clade (III), ‘LGFs’ cluster with pestiviruses in another (II), whereas ‘jingmenviruses’, orthoflaviviruses, and ‘tamanaviruses’ cluster in another (I). There is no pre-defined level or evolutionary distance range in the RdRP phylogeny (or of hallmark genes elsewhere in the ICTV taxonomy) that dictate family rank assignments. However, a threshold Jaccard distance level of 0.8–0.85 drawn through the amarilloviral grouping reproduces family rank assignments elsewhere in realm *Riboviria* (Fig. 6B).

Consequently, splitting the current family *Flaviviridae* into three families is consistent with degrees of relatedness between and within other classified RdRP-encoding RNA virus families. This split results in the removal of genera *Hepacivirus*, *Pegivirus*, and *Pestivirus* from family *Flaviviridae*, which would be restricted to genus *Orthoflavivirus* and novel genera for ‘flavi-like’ viruses.

The suggested family ‘*Pestiviridae*’ would absorb genus *Pestivirus* but also includes the multiple and highly divergent clades of ‘LGFs’. Mammalian pestiviruses possess a type IV IRES^70,71^ and major envelope proteins that are structurally homologous to those of hepaciviruses and pegiviruses^60^. However, unclassified pesti-like viruses of spiders (lineage IIm) may use cap-dependent translation and encode envelope proteins structurally unrelated to those of the originally assigned mammalian pestiviruses^60^. ‘LGFs’ are far more diverse genetically in the RdRP sequence than those of viruses in the other suggested families. However, lowering the family rank assignment threshold would create up to five or more new families of ‘pesti-like’ viruses, a step we consider inappropriate at least until these viruses are better characterized.

The remainder of flaviviruses and ‘flavi-like’ viruses group in a deep and phylogenetically well-defined third family, ‘*Hepaciviridae*’. The depth of grouping provides no support for the formation of separate families for hepaciviruses and pegiviruses. The addition of recently described ‘hepaci-like’ or ‘pegi-like’ viruses infecting fish has greatly expanded the genetic diversity of both groups and blurs the originally clear distinction between them. Generally, however, the apparent absence of a capsid-encoding sequence in pegiviruses, indicative of a likely radically different virion structure (or conceivably undetected segmentation of the pegivirus genome), differentiates pegiviruses from hepaciviruses at least for now.

The analysis provides the basis for assignments of new genera within each of the three new families (Table 1), where they are supported into genetically well-defined groups and have some commonality in phenotypic properties, such as host range, and of genome organisation. The proposed additional ten taxa, or alternatively into eight genera and three subgenera, represent obvious candidates for classification by these criteria. However, additional assignments may be made in the future pending collection of further characterised ‘flavi-like’ viruses in ongoing clinical, veterinary and entomology screens, and in metagenomic data from the widening range of invertebrate species.

Overall, we believe we have made a robust case for an evolutionarily-based reclassification of flaviviruses using an approach that puts primacy on genetic relationships of the hallmark RdRP gene. The seeming propensity of flaviviruses and potentially other RNA viruses to undergo radical changes in genome organisation such as segmentation, changes to translation mechanisms and exchanges of structural gene modules indeed reinforces the need to base classification and inference on evolutionary histories of the most stable elements within the genome. The use of protein structure relationships of RdRP and potentially other replication-associated enzymes provides an exciting new method to determine deeper evolutionary histories of RNA viruses beyond the level of family and order analysed in the current study.

## Methods

### Genome sequences

A comprehensive set of coding-complete genome sequences representing the 97 currently established flavivirus species^11^, supplemented with ‘flavi-like’ viruses with analyzed previously^60^ was used as the basic set for analysis and putative taxon assignments (Table S1; File S1). Genome regions encoding the RNA-directed RNA polymerase (RdRP) and helicase domains were extracted for amino-acid sequence deduction, alignment, and analysis.

### Sequence alignment and phylogenetic analysis

Alignment and trimming of RdRP domain amino-acids sequences with TrimAI^72^ was performed using multiple methods and conservation thresholds as previously described^60^. The phylogeny of sequences in the final alignment was reconstructed by the maximum likelihood-based IQ-TREE program version 1.6.12^73^ with an empirically determined optimal model, Lascuel + F + 10 rate categories (LG+F+R10) selected based on the minimum Bayesian information criteria (BIC) score^74^. Robustness of branching was estimated by bootstrap resampling (1,000 replicates)^75^. An unrooted tree with bootstrap support values shown for the main clades and lineages was plotted using MEGA7.0^76^.

The RdRP domain amino-acid sequence dataset was analyzed in parallel by the unweighted pair group method with arithmetic mean (UPGMA) as implemented in MEGA7.0, using Jones-Taylor-Thornton (JTT) matrix protein distances and 100 bootstrap replicates.

A temporal reconstruction of amarilloviral evolution was performed using BEAST version 10.05 using dated sequences based on sample date (or International Nucleotide Sequence Database Collaboration [INSDC] submission date if this information was not annotated; n=143), using uniform rate, constant population size, and BLOSUM protein distances as priors.

A maximum likelihood phylogenetic tree of helicase-encoding sequences was generated similarly using IQ-TREE, with the (lowest BIC) substitution model. The alignment was obtained with MAFFT^77^ and trimmed using TrimAI^72^ and the gappyout option. RdRP and helicase domain trees were compared using Tanglegram^78,79^. Potyviruses, poxvirus, and DEAH helicase family sequences were used to root the tree.

### Genome Relatedness Applied to Virus Taxonomy (GRAViTy)

Genome relatedness of flaviviruses and ‘flavi-like’ viruses to all currently classified ribovirians was performed using the GRAViTy version 2 implementation^66^ of the original algorithm^80^. Analysis was performed in a single step, using the new classification function. Default parameters were used, except for initial translated open reading frame sequence clustering inflation (6.0), translated open reading frame alignment method (G-INS-I), and protein profile hidden Markov model (PPHMM) similarity cutoff hitscore. Taxonomic assignments were bootstrapped with 10 iterations, using the sumtrees method.

### RdRP structure comparisons

We started with a previous dataset of *Flaviviridae* protein structure predictions (https://zenodo.org/records/11092288)^60^. This covers all viruses examined in the current study, with their respective polyproteins sequences broken into sequential 300 residue blocks (overlapping by 100 residues) for protein structure prediction with ColabFold and ESMFold^63,81,82^. The fragmented nature of these structures was insufficient for accurate structural alignment, therefore we queried the dataset against experimental RdRP structure references using Foldseek^64^: orthoflavivirus PDB:5F3Z, pestivirus PDB:5YF6 and hepacivirus PDB: 1C2P)^83–85^, enabling us to extract continuous RdRP domain sequences for all viruses.

Structures were predicted for these RdRP sequences using ColabFoldv1.5.5^82^, taking the highest confidence model from five predictions. These structures were filtered for average local distance difference test (lddt) prediction confidence using a semi-arbitrary cut-off of ≥80%. We also discarded structures below 400 residues in length, reasoning that heavily truncated structures may be misinformative. This yielded a final structure dataset for this study containing 400 flavivirus and ‘flavi-like’ virus RdRP structures.

RdRP structures were analysed using FoldTree to produce a structure-guided tree based on the lddt structural similarity metric, which provides a measure of structural similarity whilst accommodating for some structural flexibility^86^. In short, FoldTree^65^ performs an all-vs.-all structure comparison, driven by Foldseek^64^, to derive pairwise lddt values that are used to calculate a distance matrix and derive a Neighbour-Joining tree via QuickTree^87^. The tree was visualised and prepared for publication using iTOL^88^. Example structural models in Figure 5b were aligned for visualization using flexible FATCAT^89^, models were viewed and prepared for publication using UCSF ChimeraX^90^.

### Resource Availability

#### Lead Contact

Peter Simmonds

#### Materials availability

Not applicable, no biological materials were used in the study

#### Data and code availability

Databases, sequence alignments and raw sequence distance data provided in Suppl. Data. Correspondence and requests for materials should be addressed to PS / JCOM (RdRP sequence analysis), JG (RdRP structure analysis, RM (GRAViTy analysis) or AB (helicase analysis).

## Acknowledgements

The authors thank Anya Crane (Integrated Research Facility at Fort Detrick/National Institute of Allergy and Infectious Diseases/National Institutes of Health, Fort Detrick, Frederick, MD, USA) for critically editing the manuscript.

This work was supported in part through a Laulima Government Solutions, LLC, prime contract with the National Institute of Allergy and Infectious Diseases (Contract No. HHSN272201800013C). J.H.K. performed this work as an employee of Tunnell Government Services (TGS), a subcontractor of Laulima Government Solutions, LLC, under Contract No. HHSN272201800013C. NV acknowledges partial support from the Centers for Research in Emerging Infectious Diseases (CREID) Coordinating Research on Emerging Arboviral Threats Encompassing the NEOtropics (CREATE-NEO) 1U01AI151807 grant by the National Institutes of Health (NIH). A.B. is supported by a postdoctoral fellowship from Foundation pour la Recherche Mèdicale (grant number SPF202110014092). JG was supported by a Wellcome Trust/Royal Society Sir Henry Dale Fellowship (107653/Z/15/Z) and MRC-University of Glasgow Centre for Virus Research core support from the Medical Research Council (MC_UU_00034/1). JTS was supported by Veterans Administration Merit Review BX000207 and VA SEQCure Network grants. be interpreted as necessarily representing the official policies, either expressed or implied, of the U.S. Department of Health and Human Services or of the institutions and companies affiliated with the authors, nor does mention of trade names, commercial products, or organizations imply endorsement by the U.S. Government.

## Author contributions

PS, JHK, and members of the original ICTV Study Group (MB, JFD, AK, VL, JS and NV) conceived the study. PS, AB, JG, RM, JCOM and JHK conceptualized the experimental section. PS, AB, JG, RM and JCOM performed analyses. All authors wrote/revised the manuscript and PS supervised the work. All authors read and approved the manuscript.

## Supplementary information

The online version contains supplementary material comprising:

**Fig. S1.** Helicase domain (NS3) amino-acid sequence phylogeny supports partition of flaviviruses and ‘flavi-like’ viruses into one order and four family level clades. Fully branch-labeled version of Fig 4.

**Fig. S2.** Alignment-free hidden Markov model homology analysis supports partition of flaviviruses and ‘flavi-like’ viruses into one order and four family level clades. Fully branch-labeled version of Fig 6A.

**Fig. S3**. Alignment-free hidden Markov model homology analysis supports partition of flaviviruses and ‘flavi-like’ viruses into four main clades. Fully branch-labeled version of Fig 6B.’

**Fig. S4**. Mean pairwise amino acid sequence identities between lineages in clades I–III.

**File S1** - RdRP_tree.NWK

**File S2** - GRAViTyv2 Run parameters.json

**File S3** - RdRP_lddt_structure tree.NWK

## References

1. Simmonds, P., Becher, P., Bukh, J., Gould, E.A., Meyers, G., Monath, T., Muerhoff, S., Pletnev, A., Rico-Hesse, R., Smith, D.B., et al. (2017). ICTV virus taxonomy profile: *Flaviviridae*. J Gen Virol 98, 2–3. 10.1099/jgv.0.000672.

2. Reeves, W.C. (2001). Partners: serendipity in arbovirus research. J Vector Ecol 26, 1–6.

3. Casals, J., and Brown, L.V. (1954). Hemagglutination with arthropod-borne viruses. J Exp Med 99, 429–449. 10.1084/jem.99.5.429.

4. Fenner, F. (1976). Classification and nomenclature of viruses. Second report of the International Committee on Taxonomy of Viruses. Intervirology 7, 1–115. 10.1159/000149938.

5. Francki, R.I.B., Fauquet, C.M., Knudson, D.L., and Brown, F., eds. (1991). Classification and nomenclature of viruses. Fifth report of the International Committee on Taxonomy of Viruses. Archives of Virology Supplementum 2 (Springer-Verlag).

6. Westaway, E.G., Brinton, M.A., Gaidamovich, S.Y., Horzinek, M.C., Igarashi, A., Kääriäinen, L., Lvov, D.K., Porterfield, J.S., Russell, P.K., and Trent, D.W. (1985). *Flaviviridae*. Report of the *Togaviridae* Study Group, Vertebrate Virus Subcommittee, International Committee on Taxonomy of Viruses. Intervirology 24, 183–192. 10.1159/000149642.

7. Choo, Q.L., Richman, K.H., Han, J.H., Berger, K., Lee, C., Dong, C., Gallegos, C., Coit, D., Medina Selby, R., Barr, P.J., et al. (1991). Genetic organization and diversity of the hepatitis C virus. Proc.Natl.Acad.Sci.USA 88, 2451–2455.

8. Scheel, T.K., Simmonds, P., and Kapoor, A. (2015). Surveying the global virome: identification and characterization of HCV-related animal hepaciviruses. Antiviral Res 115, 83–93. 10.1016/j.antiviral.2014.12.014.

9. Stapleton, J.T., Foung, S., Muerhoff, A.S., Bukh, J., and Simmonds, P. (2011). The GB viruses: a review and proposed classification of GBV-A, GBV-C (HGV), and GBV-D in genus Pegivirus within the family Flaviviridae. J Gen Virol 92, 233–246. 10.1099/vir.0.027490-0.

10. Wu, Z., Wu, Y., Zhang, W., Merits, A., Simmonds, P., Wang, M., Jia, R., Zhu, D., Liu, M., Zhao, X., et al. (2020). The First Nonmammalian Pegivirus Demonstrates Efficient In Vitro Replication and High Lymphotropism. J Virol 94. 10.1128/jvi.01150-20.

11. Postler, T.S., Beer, M., Blitvich, B.J., Bukh, J., de Lamballerie, X., Drexler, J.F., Imrie, A., Kapoor, A., Karganova, G.G., Lemey, P., et al. (2023). Renaming of the genus Flavivirus to Orthoflavivirus and extension of binomial species names within the family Flaviviridae. Arch Virol 168, 224. 10.1007/s00705-023-05835-1.

12. Choo, Q.-L., Richman, K.H., Han, J.H., Berger, K., Lee, C., Dong, C., Gallegos, C., Coit, D., Medina-Selby, R., Barr, P.J., et al. (1991). Genetic organization and diversity of the hepatitis C virus. Proc Natl Acad Sci U S A 88, 2451–2455. 10.1073/pnas.88.6.2451.

13. Shi, M., Lin, X.-D., Vasilakis, N., Tian, J.-H., Li, C.-X., Chen, L.-J., Eastwood, G., Diao, X.-N., Chen, M.-H., Chen, X., et al. (2016). Divergent viruses discovered in arthropods and vertebrates revise the evolutionary history of the *Flaviviridae* and related viruses. J Virol 90, 659–669. 10.1128/JVI.02036-15.

14. Orba, Y., Matsuno, K., Nakao, R., Kryukov, K., Saito, Y., Kawamori, F., Loza Vega, A., Watanabe, T., Maemura, T., Sasaki, M., et al. (2021). Diverse mosquito-specific flaviviruses in the Bolivian Amazon basin. J Gen Virol 102. 10.1099/jgv.0.001518.

15. Blasdell, K.R., Wynne, J.W., Perera, D., and Firth, C. (2021). First detection of a novel ‘unknown host’ flavivirus in a Malaysian rodent. Access Microbiol 3, 000223. 10.1099/acmi.0.000223.

16. Gravina, H.D., Suzukawa, A.A., Zanluca, C., Cardozo Segovia, F.M., Tschá, M.K., Martins da Silva, A., Faoro, H., da Silva Ribeiro, R., Mendoza Torres, L.P., Rojas, A., et al. (2019). Identification of insect-specific flaviviruses in areas of Brazil and Paraguay experiencing endemic arbovirus transmission and the description of a novel flavivirus infecting Sabethes belisarioi. Virology 527, 98–106. 10.1016/j.virol.2018.11.008.

17. Kobayashi, D., Watanabe, M., Faizah, A.N., Amoa-Bosompem, M., Higa, Y., Tsuda, Y., Sawabe, K., and Isawa, H. (2021). Discovery of a Novel Flavivirus (Flaviviridae) From the Horse Fly, Tabanus rufidens (Diptera: Tabanidae): The Possible Coevolutionary Relationships Between the Classical Insect-Specific Flaviviruses and Host Dipteran Insects. J Med Entomol 58, 880–890. 10.1093/jme/tjaa193.

18. Korkusol, A., Takhampunya, R., Hang, J., Jarman, R.G., Tippayachai, B., Kim, H.C., Chong, S.T., Davidson, S.A., and Klein, T.A. (2017). A novel flavivirus detected in two Aedes spp. collected near the demilitarized zone of the Republic of Korea. J Gen Virol 98, 1122–1131. 10.1099/jgv.0.000746.

19. McLean, B.J., Hobson-Peters, J., Webb, C.E., Watterson, D., Prow, N.A., Nguyen, H.D., Hall-Mendelin, S., Warrilow, D., Johansen, C.A., Jansen, C.C., et al. (2015). A novel insect-specific flavivirus replicates only in Aedes-derived cells and persists at high prevalence in wild Aedes vigilax populations in Sydney, Australia. Virology 486, 272–283. 10.1016/j.virol.2015.07.021.

20. Smith, D.B., Meyers, G., Bukh, J., Gould, E.A., Monath, T., Scott Muerhoff, A., Pletnev, A., Rico-Hesse, R., Stapleton, J.T., Simmonds, P., and Becher, P. (2017). Proposed revision to the taxonomy of the genus Pestivirus, family Flaviviridae. J Gen Virol 98, 2106–2112. 10.1099/jgv.0.000873.

21. Smith, D.B., Becher, P., Bukh, J., Gould, E.A., Meyers, G., Monath, T., Muerhoff, A.S., Pletnev, A., Rico-Hesse, R., Stapleton, J.T., and Simmonds, P. (2016). Proposed update to the taxonomy of the genera Hepacivirus and Pegivirus within the Flaviviridae family. J Gen Virol 97, 2894–2907. 10.1099/jgv.0.000612.

22. Hartlage, A.S., Cullen, J.M., and Kapoor, A. (2016). The Strange, Expanding World of Animal Hepaciviruses. Annual review of virology 3, 53–75. 10.1146/annurev-virology-100114-055104.

23. Ridpath, J.F. (2013). A need to define characteristics to be used in the taxonomy of the expanding pestivirus genus. Berl Munch Tierarztl Wochenschr 126, 462–467. 10.2376/0005-9366-126-462.

24. Shi, M., Lin, X.D., Vasilakis, N., Tian, J.H., Li, C.X., Chen, L.J., Eastwood, G., Diao, X.N., Chen, M.H., Chen, X., et al. (2015). Divergent Viruses Discovered in Arthropods and Vertebrates Revise the Evolutionary History of the Flaviviridae and Related Viruses. J Virol 90, 659–669. 10.1128/jvi.02036-15.

25. Matsumura, E.E., Nerva, L., Nigg, J.C., Falk, B.W., and Nouri, S. (2016). Complete Genome Sequence of the Largest Known Flavi-Like Virus, Diaphorina citri flavi-like virus, a Novel Virus of the Asian Citrus Psyllid, Diaphorina citri. Genome Announc 4. 10.1128/genomeA.00946-16.

26. Kartashov, M.Y., Gladysheva, A.V., Shvalov, A.N., Tupota, N.L., Chernikova, A.A., Ternovoi, V.A., and Loktev, V.B. (2023). Novel Flavi-like virus in ixodid ticks and patients in Russia. Ticks Tick Borne Dis 14, 102101. 10.1016/j.ttbdis.2022.102101.

27. Petrone, M.E., Grove, J., Mélade, J., Mifsud, J.C.O., Parry, R.H., Marzinelli, E.M., and Holmes, E.C. (2024). A ∼40-kb flavi-like virus does not encode a known error-correcting mechanism. Proc Natl Acad Sci U S A 121, e2403805121. 10.1073/pnas.2403805121.

28. Qin, X.-C., Shi, M., Tian, J.-H., Lin, X.-D., Gao, D.-Y., He, J.-R., Wang, J.-B., Li, C.-X., Kang, Y.-J., Yu, B., et al. (2014). A tick-borne segmented RNA virus contains genome segments derived from unsegmented viral ancestors. Proc Natl Acad Sci U S A 111, 6744–6749. 10.1073/pnas.1324194111.

29. Ladner, J.T., Wiley, M.R., Beitzel, B., Auguste, A.J., Dupuis, A.P., 2nd, Lindquist, M.E., Sibley, S.D., Kota, K.P., Fetterer, D., Eastwood, G., et al. (2016). A multicomponent animal virus isolated from mosquitoes. Cell host & microbe 20, 357–367. 10.1016/j.chom.2016.07.011.

30. Zhang, S., Yang, C., Qiu, Y., Liao, R., Xuan, Z., Ren, F., Dong, Y., Xie, X., Han, Y., Wu, D., et al. (2024). Conserved untranslated regions of multipartite viruses: Natural markers of novel viral genomic components and tags of viral evolution. Virus evolution 10, veae004. 10.1093/ve/veae004.

31. Paraskevopoulou, S., Käfer, S., Zirkel, F., Donath, A., Petersen, M., Liu, S., Zhou, X., Drosten, C., Misof, B., and Junglen, S. (2021). Viromics of extant insect orders unveil the evolution of the flavi-like superfamily. Virus evolution 7, veab030. 10.1093/ve/veab030.

32. Colmant, A.M.G., Charrel, R.N., and Coutard, B. (2022). Jingmenviruses: Ubiquitous, understudied, segmented flavi-like viruses. Frontiers in microbiology 13, 997058. 10.3389/fmicb.2022.997058.

33. Mifsud, J.C.O., Costa, V.A., Petrone, M.E., Marzinelli, E.M., Holmes, E.C., and Harvey, E. (2023). Transcriptome mining extends the host range of the Flaviviridae to non-bilaterians. Virus evolution 9, veac124. 10.1093/ve/veac124.

34. Parry, R., and Asgari, S. (2019). Discovery of novel crustacean and cephalopod flaviviruses: insights into the evolution and circulation of flaviviruses between marine invertebrate and vertebrate hosts. J Virol 93, e00432–00419. 10.1128/JVI.00432-19.

35. Lay, C.L., Shi, M., Buček, A., Bourguignon, T., Lo, N., and Holmes, E.C. (2020). Unmapped RNA Virus Diversity in Termites and their Symbionts. Viruses 12. 10.3390/v12101145.

36. Dong, X., Wang, G., Hu, T., Li, J., Li, C., Cao, Z., Shi, M., Wang, Y., Zou, P., Song, J., et al. (2021). A Novel Virus of Flaviviridae Associated with Sexual Precocity in Macrobrachium rosenbergii. mSystems 6, e0000321. 10.1128/mSystems.00003-21.

37. Bekal, S., Domier, L.L., Gonfa, B., McCoppin, N.K., Lambert, K.N., and Bhalerao, K. (2014). A novel flavivirus in the soybean cyst nematode. J Gen Virol 95, 1272–1280. 10.1099/vir.0.060889-0.

38. Dheilly, N.M., Lucas, P., Blanchard, Y., and Rosario, K. (2022). A World of Viruses Nested within Parasites: Unraveling Viral Diversity within Parasitic Flatworms (Platyhelminthes). Microbiol Spectr 10, e0013822. 10.1128/spectrum.00138-22.

39. Chen, Y.-M., Sadiq, S., Tian, J.-H., Chen, X., Lin, X.-D., Shen, J.-J., Chen, H., Hao, Z.-Y., Wille, M., Zhou, Z.-C., et al. (2022). RNA viromes from terrestrial sites across China expand environmental viral diversity. Nat Microbiol 7, 1312–1323. 10.1038/s41564-022-01180-2.

40. Hewson, I., Johnson, M.R., and Tibbetts, I.R. (2020). An Unconventional Flavivirus and Other RNA Viruses in the Sea Cucumber (Holothuroidea; Echinodermata) Virome. Viruses 12. 10.3390/v12091057.

41. Geoghegan, J.L., Di Giallonardo, F., Cousins, K., Shi, M., Williamson, J.E., and Holmes, E.C. (2018). Hidden diversity and evolution of viruses in market fish. Virus evolution 4, vey031. 10.1093/ve/vey031.

42. Skoge, R.H., Brattespe, J., Økland, A.L., Plarre, H., and Nylund, A. (2018). New virus of the family Flaviviridae detected in lumpfish (Cyclopterus lumpus). Arch Virol 163, 679–685. 10.1007/s00705-017-3643-3.

43. Soto, E., Camus, A., Yun, S., Kurobe, T., Leary, J.H., Rosser, T.G., Dill-Okubo, J.A., Nyaoke, A.C., Adkison, M., Renger, A., and Ng, T.F.F. (2020). First Isolation of a Novel Aquatic Flavivirus from Chinook Salmon (Oncorhynchus tshawytscha) and Its In Vivo Replication in a Piscine Animal Model. J Virol 94. 10.1128/jvi.00337-20.

44. Costa, V.A., Mifsud, J.C.O., Gilligan, D., Williamson, J.E., Holmes, E.C., and Geoghegan, J.L. (2021). Metagenomic sequencing reveals a lack of virus exchange between native and invasive freshwater fish across the Murray-Darling Basin, Australia. Virus evolution 7, veab034. 10.1093/ve/veab034.

45. Parry, R.H., Slonchak, A., Campbell, L.J., Newton, N.D., Debat, H.J., Gifford, R.J., and Khromykh, A.A. (2023). A novel tamanavirus (Flaviviridae) of the European common frog (Rana temporaria) from the UK. J Gen Virol 104. 10.1099/jgv.0.001927.

46. Urayama, S.-I., Takaki, Y., and Nunoura, T. (2016). FLDS: A Comprehensive dsRNA Sequencing Method for Intracellular RNA Virus Surveillance. Microbes Environ 31, 33–40. 10.1264/jsme2.ME15171.

47. Forgia, M., Daghino, S., Chiapello, M., Ciuffo, M., and Turina, M. (2024). New clades of viruses infecting the obligatory biotroph *Bremia lactucae* representing distinct evolutionary trajectory for viruses infecting oomycetes. Virus evolution 10, veae003. 10.1093/ve/veae003.

48. Debat, H., and Bejerman, N. (2023). Two novel flavi-like viruses shed light on the plant-infecting koshoviruses. Arch Virol 168, 184. 10.1007/s00705-023-05813-7.

49. Schönegger, D., Marais, A., Faure, C., and Candresse, T. (2022). A new flavi-like virus identified in populations of wild carrots. Arch Virol 167, 2407–2409. 10.1007/s00705-022-05544-1.

50. Bamford, C.G.G., de Souza, W.M., Parry, R., and Gifford, R.J. (2022). Comparative analysis of genome-encoded viral sequences reveals the evolutionary history of flavivirids (family Flaviviridae). Virus evolution 8, veac085. 10.1093/ve/veac085.

51. Simmonds, P., Adriaenssens, E.M., Zerbini, F.M., Abrescia, N.G.A., Aiewsakun, P., Alfenas-Zerbini, P., Bao, Y., Barylski, J., Drosten, C., Duffy, S., et al. (2023). Four principles to establish a universal virus taxonomy. PLoS biology 21, e3001922. 10.1371/journal.pbio.3001922.

52. Nasir, A., Romero-Severson, E., and Claverie, J.M. (2020). Investigating the Concept and Origin of Viruses. Trends Microbiol 28, 959–967. 10.1016/j.tim.2020.08.003.

53. Brussow, H. (2009). The not so universal tree of life or the place of viruses in the living world. Philosophical transactions of the Royal Society of London. Series B, Biological sciences 364, 2263–2274. 10.1098/rstb.2009.0036.

54. Krupovic, M., Dolja, V.V., and Koonin, E.V. (2019). Origin of viruses: primordial replicators recruiting capsids from hosts. Nat Rev Microbiol 17, 449–458. 10.1038/s41579-019-0205-6.

55. Kuhn, J.H., Wolf, Y.I., Krupovic, M., Zhang, Y.Z., Maes, P., Dolja, V.V., and Koonin, E.V. (2019). Classify viruses - the gain is worth the pain. Nature 566, 318–320. 10.1038/d41586-019-00599-8.

56. Koonin, E.V., Dolja, V.V., Krupovic, M., Varsani, A., Wolf, Y.I., Yutin, N., Zerbini, F.M., and Kuhn, J.H. (2020). Global Organization and Proposed Megataxonomy of the Virus World. Microbiol Mol Biol Rev 84. 10.1128/mmbr.00061-19.

57. Koonin, E.V., Kuhn, J.H., Dolja, V.V., and Krupovic, M. (2024). Megataxonomy and global ecology of the virosphere. The ISME journal 18. 10.1093/ismejo/wrad042.

58. Belsham, G.J. (2009). Divergent picornavirus IRES elements. Virus Res 139, 183–192. 10.1016/j.virusres.2008.07.001.

59. Arhab, Y., Bulakhov, A.G., Pestova, T.V., and Hellen, C.U.T. (2020). Dissemination of Internal Ribosomal Entry Sites (IRES) Between Viruses by Horizontal Gene Transfer. Viruses 12. 10.3390/v12060612.

60. Mifsud, J.C.O., Lytras, S., Oliver, M.R., Toon, K., Costa, V.A., Holmes, E.C., and Grove, J. (2024). Mapping glycoprotein structure reveals Flaviviridae evolutionary history. Nature. 10.1038/s41586-024-07899-8.

61. Vaney, M.C., Dellarole, M., Duquerroy, S., Medits, I., Tsouchnikas, G., Rouvinski, A., England, P., Stiasny, K., Heinz, F.X., and Rey, F.A. (2022). Evolution and activation mechanism of the flavivirus class II membrane-fusion machinery. Nat Commun 13, 3718. 10.1038/s41467-022-31111-y.

62. Jumper, J., Evans, R., Pritzel, A., Green, T., Figurnov, M., Ronneberger, O., Tunyasuvunakool, K., Bates, R., Žídek, A., Potapenko, A., et al. (2021). Highly accurate protein structure prediction with AlphaFold. Nature 596, 583–589. 10.1038/s41586-021-03819-2.

63. Varadi, M., Bertoni, D., Magana, P., Paramval, U., Pidruchna, I., Radhakrishnan, M., Tsenkov, M., Nair, S., Mirdita, M., Yeo, J., et al. (2024). AlphaFold Protein Structure Database in 2024: providing structure coverage for over 214 million protein sequences. Nucleic Acids Res 52, D368–d375. 10.1093/nar/gkad1011.

64. van Kempen, M., Kim, S.S., Tumescheit, C., Mirdita, M., Lee, J., Gilchrist, C.L.M., Söding, J., and Steinegger, M. (2024). Fast and accurate protein structure search with Foldseek. Nat Biotechnol 42, 243–246. 10.1038/s41587-023-01773-0.

65. Moi, D., Bernard, C., Steinegger, M., Nevers, Y., Langleib, M., and Dessimoz, C. (2023). Structural phylogenetics unravels the evolutionary diversification of communication systems in gram-positive bacteria and their viruses. bioRxiv.

66. Mayne, R., Aiewsakun, P., Turner, D., Adriaenssens, E.M., and Simmonds, P. (2024). GRAViTy-V2: a grounded viral taxonomy application. bioRxiv, 2024.2007.2026.605250. 10.1101/2024.07.26.605250.

67. Marin, M.S., Zanotto, P.M., Gritsun, T.S., and Gould, E.A. (1995). Phylogeny of TYU, SRE, and CFA virus: different evolutionary rates in the genus Flavivirus. Virology 206, 1133–1139. 10.1006/viro.1995.1038.

68. Kobayashi, K., Atsumi, G., Iwadate, Y., Tomita, R., Chiba, K., Akasaka, S., Nishihara, M., Takahashi, H., Yamaoka, N., Nishiguchi, M., and Sekine, K.T. (2013). Gentian Kobu-sho-associated virus: a tentative, novel double-stranded RNA virus that is relevant to gentian Kobu-sho syndrome. J Gen Plant Pathol 79, 56–63. 10.1007/s10327-012-0423-5.

69. Wolf, Y.I., Kazlauskas, D., Iranzo, J., Lucía-Sanz, A., Kuhn, J.H., Krupovic, M., Dolja, V.V., and Koonin, E.V. (2018). Origins and Evolution of the Global RNA Virome. mBio 9. 10.1128/mBio.02329-18.

70. Fletcher, S.P., and Jackson, R.J. (2002). Pestivirus internal ribosome entry site (IRES) structure and function: elements in the 5’ untranslated region important for IRES function. J Virol 76, 5024–5033. 10.1128/jvi.76.10.5024-5033.2002.

71. Reusken, C.B.E.M., Dalebout, T.J., Eerligh, P., Bredenbeek, P.J., and Spaan, W.J.M. (2003). Analysis of hepatitis C virus/classical swine fever virus chimeric 5’NTRs: sequences within the hepatitis C virus IRES are required for viral RNA replication. J Gen Virol 84, 1761–1769. 10.1099/vir.0.19063-0.

72. Capella-Gutiérrez, S., Silla-Martínez, J.M., and Gabaldón, T. (2009). trimAl: a tool for automated alignment trimming in large-scale phylogenetic analyses. Bioinformatics (Oxford, England) 25, 1972–1973. 10.1093/bioinformatics/btp348.

73. Nguyen, L.-T., Schmidt, H.A., von Haeseler, A., and Minh, B.Q. (2015). IQ-TREE: a fast and effective stochastic algorithm for estimating maximum-likelihood phylogenies. Mol Biol Evol 32, 268–274. 10.1093/molbev/msu300.

74. Kalyaanamoorthy, S., Minh, B.Q., Wong, T.K.F., von Haeseler, A., and Jermiin, L.S. (2017). ModelFinder: fast model selection for accurate phylogenetic estimates. Nature methods 14, 587–589. 10.1038/nmeth.4285.

75. Hoang, D.T., Chernomor, O., von Haeseler, A., Minh, B.Q., and Vinh, L.S. (2018). UFBoot2: Improving the Ultrafast Bootstrap Approximation. Mol Biol Evol 35, 518–522. 10.1093/molbev/msx281.

76. Kumar, S., Stecher, G., and Tamura, K. (2016). MEGA7: Molecular Evolutionary Genetics Analysis Version 7.0 for Bigger Datasets. Mol Biol Evol 33, 1870–1874. 10.1093/molbev/msw054.

77. Katoh, K., and Standley, D.M. (2013). MAFFT multiple sequence alignment software version 7: improvements in performance and usability. Mol Biol Evol 30, 772–780. 10.1093/molbev/mst010.

78. Scornavacca, C., Zickmann, F., and Huson, D.H. (2011). Tanglegrams for rooted phylogenetic trees and networks. Bioinformatics (Oxford, England) 27, i248–256. 10.1093/bioinformatics/btr210.

79. Galili, T. (2015). dendextend: an R package for visualizing, adjusting and comparing trees of hierarchical clustering. Bioinformatics (Oxford, England) 31, 3718–3720. 10.1093/bioinformatics/btv428.

80. Aiewsakun, P., and Simmonds, P. (2018). The genomic underpinnings of eukaryotic virus taxonomy: creating a sequence-based framework for family-level virus classification. Microbiome 6, 38. 10.1186/s40168-018-0422-7.

81. Lin, Z., Akin, H., Rao, R., Hie, B., Zhu, Z., Lu, W., Smetanin, N., Verkuil, R., Kabeli, O., Shmueli, Y., et al. (2023). Evolutionary-scale prediction of atomic-level protein structure with a language model. Science 379, 1123–1130. 10.1126/science.ade2574.

82. Mirdita, M., Schütze, K., Moriwaki, Y., Heo, L., Ovchinnikov, S., and Steinegger, M. (2022). ColabFold: making protein folding accessible to all. Nature methods 19, 679–682. 10.1038/s41592-022-01488-1.

83. Liu, W., Shi, X., and Gong, P. (2018). A unique intra-molecular fidelity-modulating mechanism identified in a viral RNA-dependent RNA polymerase. Nucleic Acids Res 46, 10840–10854. 10.1093/nar/gky848.

84. Noble, C.G., Lim, S.P., Arora, R., Yokokawa, F., Nilar, S., Seh, C.C., Wright, S.K., Benson, T.E., Smith, P.W., and Shi, P.Y. (2016). A Conserved Pocket in the Dengue Virus Polymerase Identified through Fragment-based Screening. J Biol Chem 291, 8541–8548. 10.1074/jbc.M115.710731.

85. Lesburg, C.A., Cable, M.B., Ferrari, E., Hong, Z., Mannarino, A.F., and Weber, P.C. (1999). Crystal structure of the RNA-dependent RNA polymerase from hepatitis C virus reveals a fully encircled active site. Nat.Struct.Biol. 6, 937–943.

86. Mariani, V., Biasini, M., Barbato, A., and Schwede, T. (2013). lDDT: a local superposition-free score for comparing protein structures and models using distance difference tests. Bioinformatics (Oxford, England) 29, 2722–2728. 10.1093/bioinformatics/btt473.

87. Howe, K., Bateman, A., and Durbin, R. (2002). QuickTree: building huge Neighbour-Joining trees of protein sequences. Bioinformatics (Oxford, England) 18, 1546–1547. 10.1093/bioinformatics/18.11.1546.

88. Letunic, I., and Bork, P. (2024). Interactive Tree of Life (iTOL) v6: recent updates to the phylogenetic tree display and annotation tool. Nucleic Acids Res 52, W78–w82. 10.1093/nar/gkae268.

89. Bittrich, S., Segura, J., Duarte, J.M., Burley, S.K., and Rose, Y. (2024). RCSB protein Data Bank: exploring protein 3D similarities via comprehensive structural alignments. Bioinformatics (Oxford, England) 40. 10.1093/bioinformatics/btae370.

90. Meng, E.C., Goddard, T.D., Pettersen, E.F., Couch, G.S., Pearson, Z.J., Morris, J.H., and Ferrin, T.E. (2023). UCSF ChimeraX: Tools for structure building and analysis. Protein Sci 32, e4792. 10.1002/pro.4792.

91. Blum, M., Andreeva, A., Florentino, L.C., Chuguransky, S.R., Grego, T., Hobbs, E., Pinto, B.L., Orr, A., Paysan-Lafosse, T., Ponamareva, I., et al. (2024). InterPro: the protein hsequence classification resource in 2025. Nucleic Acids Res. 10.1093/nar/gkae1082.

